# Clots reveal anomalous elastic behavior of fiber networks

**DOI:** 10.1101/2023.03.24.534185

**Authors:** Andrei Zakharov, Myra Awan, Terrence Cheng, Arvind Gopinath, Sang-Joon John Lee, Anand K. Ramasubramanian, Kinjal Dasbiswas

## Abstract

The mechanical properties of many soft natural and synthetic biological materials are relevant to their function. The emergence of these properties from the collective response of the structural components of the material to external stress as well as to intrinsic cell traction, remains poorly understood. Here, we examine the nonlinear elastic behavior of blood clots by combining microscopy and rheological measurements with an elastic network model that accounts for the stretching, bending, and buckling of constituent fibrin fibers. We show that the inhibition of fibrin crosslinking reduces fiber bending stiffness and introduces an atypical fiber buckling-induced softening regime at intermediate shear, before the well-characterized stiffening regime. We also show that crosslinking and platelet contraction significantly alter force propagation in the network in a strain-dependent manner. Our mechanics-based model, supported by experiments, provides a framework to understand the origins of characteristic and anomalous regimes of non-linear elastic response not only in blood clots, but also more generally in active biopolymer networks.

## INTRODUCTION

Fibrous materials form the structural and functional basis for numerous biological processes and biomedical applications (*1*). Many fibrous biomaterials including actin, collagen, and fibrin occur as networks endowed with unique, non-linear mechanical properties that are key to their biological functions such as maintaining structural integrity, tissue architecture, and facilitating cell-cell communication (*2*). Of significance, the branched network of fibrin fibers is the fundamental building block of blood clots, and is integral to biomedical applications such as tissue scaffolding and surgical adhesives (*3*). Fibrin networks provide optimal strength, stiffness, and stability appropriate for these physiological processes (*4*). A fibrin network results from the thrombin-catalyzed polymerization of fibrinogen monomers into protofibrils, which are crosslinked by the transglutaminase enzyme FXIII-A to form the complex hierarchical structure of fibrin fibers (*1*). The network is strengthened by the forces generated by the active contraction of platelets bound to fibrin, thus prestressing the network (*5*); and it is also modified to a lesser, nonetheless important, extent by the passive inclusions of red blood cells (RBCs) (*6*). Although the mechanics of fibrin networks has been extensively studied (*7*), the ability to predict the non-linear mechanical behavior of the network from the molecular-scale structure and topology of constituent fibers remains elusive.

The mechanical response of blood clots to external loads is a combined response of the fibrin network, platelets associated with the network, and of the void spaces composed of plasma and the RBCs. To understand the overall structure-mechanics relationships of the composite blood clot, several biomechanical models of varying scope have been developed (recently reviewed in (*8*)). These models include constitutive or phenomenological approaches to study mechanical properties of clots as a function of clot composition (*9*),(*10*); continuum models to predict the macroscopic, large deformation of clots such as for predicting viscoelastic responses and clot rupture (*11*),(*12*); mesoscopic models that explicitly account for the mechanics of individual fibers as a collection of elastic elements connected to form two-dimensional (2D) or three-dimensional (3D) networks with prescribed topology (*13*),(*14*),(*15*); and microscale, molecular models of the unfolding of single fibrin fibers (*16*). Of relevance to the current work, mesoscopic models not only describe the elastic response of the networks under macroscopic deformation modes such as shear or uniaxial tension but also provide important insights into local fiber deformation, and long-range force transmission due to platelet-fiber interactions (*17*),(*18*).

In this work, we aim to understand the impact of fiber crosslinking and platelet-mediated network contraction on local and macroscopic clot mechanics. Experiments have revealed that crosslinking increases clot compaction, confers mechanical stability, and also enhances chemical stability by protecting against fibrinolysis (*19*),(*20*),(*21*). Atomic force microscopy (AFM) and optical tweezer studies have shown that these crosslinks significantly increase the extensibility and elasticity of even single fibrin fibers (*22*),(*23*). In addition, it has been shown that anisotropic, fibrous materials, such as collagen and fibrin, transmit forces over a longer range compared with linear elastic materials (*24*),(*25*). Such mechanical signaling can be independent of or act complementary to chemical signaling, providing a longer-ranged and faster pathway for cell-cell communications in tissues (*26*),(*27*). However, the effect of crosslinking in the stress-bearing elements, i.e., in individual fibrin fibers, on the mechanics of 3D fibrin networks, is poorly understood. Specifically, the manner in which the fibrin network serves as a conduit for the transmission of contractile forces generated by the platelets, and the role played by fibrin crosslinking in this process, remains unknown (*28*),(*29*).

To this end, we sought to understand the physical forces governing the mechanical response of clots to externally imposed shear loading. Our experiments reveal a heretofore unreported, anomalous, regime of shear softening in clots, characterized by an unexpected dip in the shear modulus with increasing shear stress, when fibrin crosslinking is inhibited. Importantly this softening occurs in an important biophysical regime of shear, similar to stresses induced by blood flow (*30*). Motivated by these observations, we develop a minimal elastic network model to quantitatively study clot mechanics, and examine the network response to applied shear. To describe the typical structure of a fibrin network (green) with embedded platelet aggregates (red) shown in the experimental image in **Fig. 1A**, we create a two-dimensional (2D) elastic network model comprising fibers represented by bonds in an irregular triangular network. The mechanical response of a 2D planar network is similar to, and captures the stiffening behavior of a 3D network as well (*31*). Disorder in the network is introduced by removing bonds randomly with a probability *1-p*, to reach an average coordination number. The average coordination number 〈*z*〉 is expected to be between 3 and 4 for fiber networks, these values correspond to branching and crossing fibers, respectively.

**Fig. 1.**
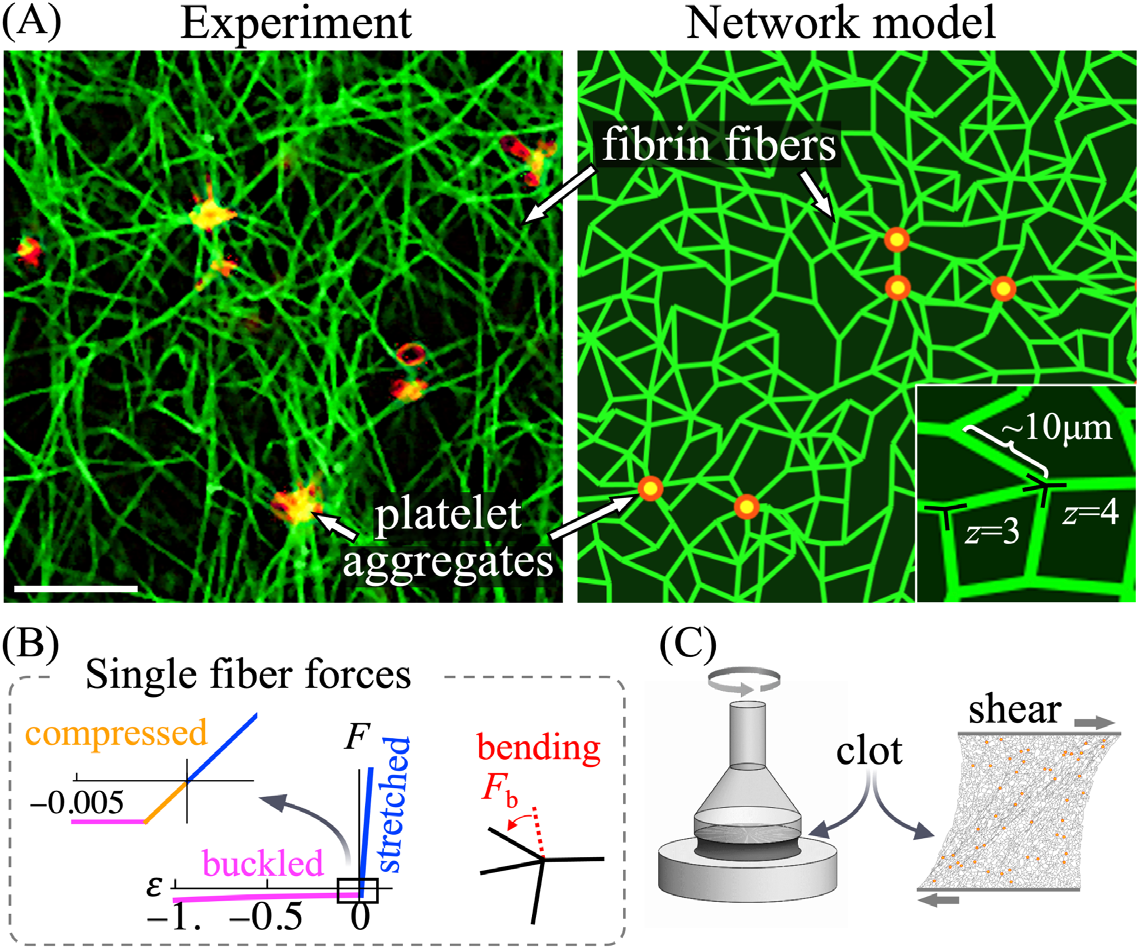
Active network model of plasma clots. **(A)** Microscopic image of platelet rich plasma (PRP) clot and its approximation by the 2D elastic network model of similar connectivity and length scale (scale bar: 10 μm). Fibrin fibers (green) forming a network are modeled by randomly oriented bonds connected at nodes representing branch points. Platelets (red) are modeled as point sources of contractile forces. The network topology is determined by the number of bonds connected at each node (the average coordination number 〈*z*〉) and by the spacing between the nodes (here, the average value is chosen to be 10 μm, similar to the average fibrin length observed in microscope images). **(B)** The reaction force *F* of each fiber to an applied axial strain *ε* exhibits three distinct regimes: stiff linear stretching, compression, and soft buckling when axial compression exceeds a small critical value (inset). In addition, a transverse load acting on a fiber is resisted by bending forces (*F_b_*) at each node when the angle between any two neighboring bonds deviates from its reference value. **(C)** Shear strain was applied to plasma clots in rheometry experiments, and in model elastic network simulations, to test their elastic response to deformation.

The elastic response of a fiber network to external shear depends on both individual fiber mechanics as well as the network architecture (*32*). Although the network is sheared at a macroscopic scale, individual fibers may stretch, compress, bend, and buckle under imposed forces, where the typical constitutive relation for a fiber is depicted in Fig. 1B. Importantly the network stability is governed by the rigidity percolation threshold, which for a network of central force springs (equal and opposite forces along the spring on a connected pair of nodes) in 2D is predicted from constraint counting originally due to Maxwell (*33*) to be 〈*z*〉 = 4. Below this isostatic point for central force springs, a fiber network can be stabilized by the bending rigidity of fibers (*34*). Since the characteristic bending energy of slender fibers is much lower than stretching energy (*35*) networks in the under-coordinated regime (〈*z*〉 < 4) respond to external shear through floppy, bending-dominated deformation modes. These floppy modes comprise rotating bonds resulting in an anomalous elastic regime not described by a continuum elastic theory (*36*). At higher shear, bonds are aligned in the shear direction and such floppy modes are no longer available. As a result the network exhibits a stiffening transition from a bending to a stretching-dominated regime (*37*). Such a bending-to-stretching stiffening transition has been shown to occur in collagen (*31*), another structured fibrous material, and is expected to be a key contributor to the nonlinear shear stiffening of biopolymer networks in general.

In this work, we combine experimental measurements with detailed numerical simulations to investigate the non-linear mechanics of sheared blood clots. We numerically compute the mechanical equilibrium states of the model elastic networks under external shear, as detailed in the *Methods* section. The model results are then compared with experiments that subject plasma clots to shear in a rheometer (Fig. 1C), to infer mechanical properties of the fibrin network. Our model predicts an abrupt clot stiffening associated with transition from bending to stretching dominated regime when strain increases, and consistent with experimental observations, it also predicts a softening dip in the shear modulus, when the bending stiffness of individual fibers is reduced. We quantitatively analyze the competition between elastic energy contributions from different fiber deformation modes that enable us to elucidate the different observed regimes in the macroscopic mechanical response of the clot. Our model also reveals that the mechanical force transmission through the network is strongly influenced by crosslinking, and provides insights into the importance of crosslinking on the prestress induced by platelets that contracts and stiffens clots.

## RESULTS

### Crosslinking alters the shear response of fibrin networks by enhancing bending stiffness

To examine the mechanical properties of fibrin networks derived from blood clots, we performed rheological experiments on plasma devoid of red blood cells (RBCs). We thus avoid the possible complicating modifications that RBCs make to the structure and mechanics of clots, such as enhanced viscosity (*6*) and compressive stiffening (*38*). To further isolate the contribution of platelets, we first prepared platelet-poor plasma (PPP) clots. The “clot stabilizing factor” FXIII-A crosslinks (or ligates) protofibrils within fibrin fibers by introducing covalent linkages between the *α*-chains and *γ*-chains of fibrin monomers. The *γ-γ* crosslinks are formed between adjacent fibrinogen monomers within a protofibril, while the *α-α* crosslinks ligate across two protofibril strands. To understand the importance of protofibril crosslinking to clot structure and mechanics, we also inhibited the crosslinker FXIII-A by treatment with the small molecule inhibitor T101 (*20*). The resulting structural differences between PPP clots with and without crosslinking are shown in Fig. 2A. The inhibition of the crosslinker FXIII-A by T101 degenerates inter-protofibril lateral *α-α* interactions, and also weakens intra-protofibril end-to-end *γ-γ* interactions, in a dose-dependent manner (*39*). The inhibition of these crosslinks does not affect the formation of protofibrils, but reduces the rates of their lateral aggregation and branching. On purely mechanical grounds, we expect that reduced lateral links between protofibrils will result in easier bending of the composite fibers they constitute (*40*), as illustrated in Fig. 2B. Consistent with this expectation, more bent and buckled fiber shapes, including kinks, are observed in **Fig. 2A** (right panel) when crosslinking is inhibited by T101 treatment. Sharp bends along with wavy and crimped structures of uncrosslinked fibrin are clearly visible in the inset of Fig. 2A.

**Fig. 2.**
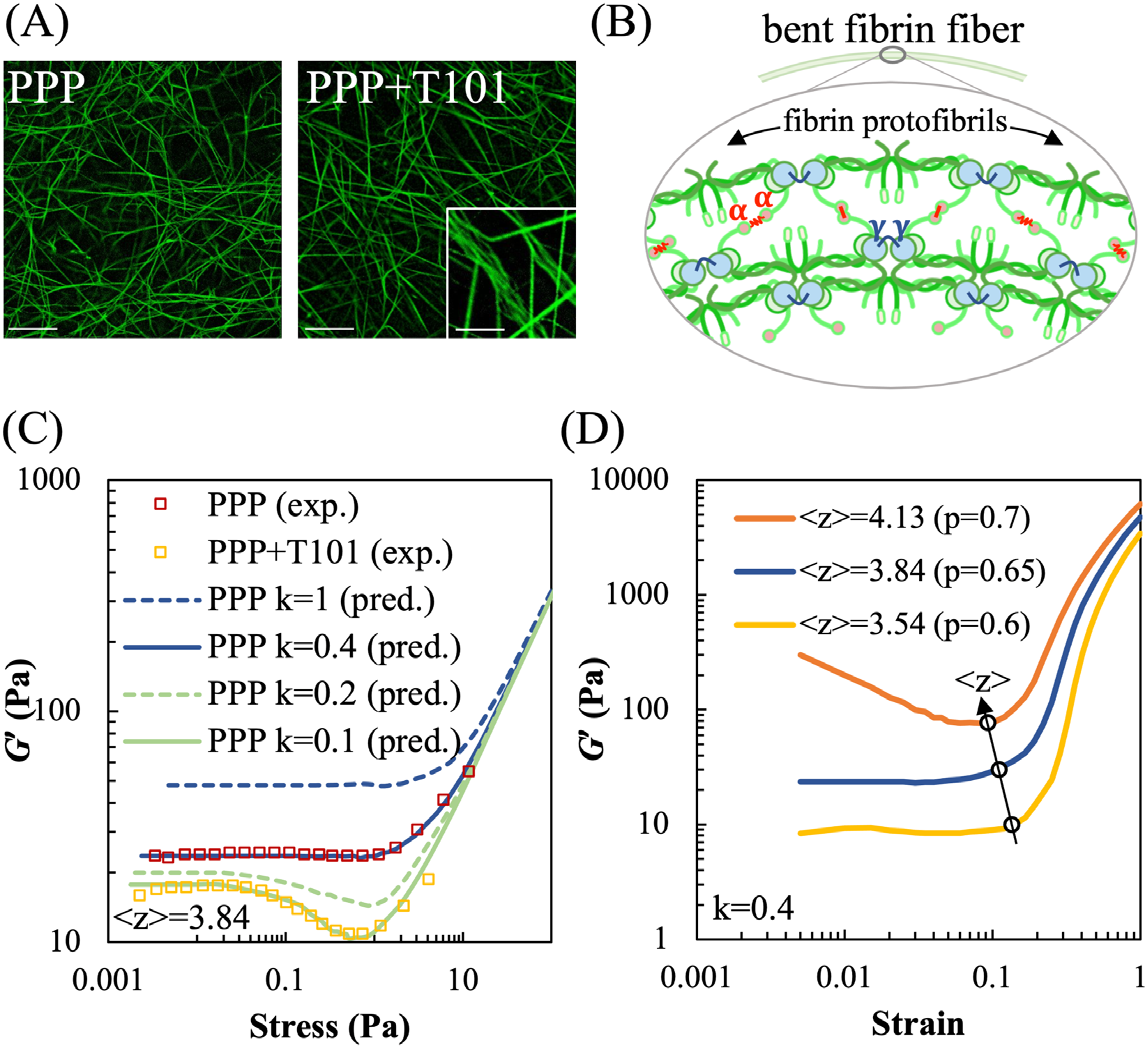
Effect of fibrin crosslinking on shear stiffness regimes of passive platelet poor plasma (PPP) clots. **(A)** Microscopic images of crosslinked (PPP) and uncrosslinked (PPP+T101) clots. Inset on the right shows a loosely crosslinked and crimped protofibrils due to inhibition of FXIII-A by T101 treatment. Scale bars: 10 μm (5 μm for inset). **(B)** Schematic of bent fibrin fiber showing protofibril structure. Fibrin oligomers assemble into protofibrils, which, in turn, are crosslinked by *γ-γ*, *α-α*, and *α-γ* interactions by FXIII-A. The crosslinks are expected to enhance bending stiffness of fibers. **(C)** Shear modulus (*G’*) of crosslinked (PPP) and uncrosslinked (PPP+T101) clots show a highly non-linear dependence on applied stress in both experiment (squares) and simulations (lines). The factor *k* and 〈*z*〉 in the simulated curves represent the reduced bending modulus from a reference value, and average coordination number of network nodes, respectively. Simulations were performed at varying values of the parameters *k* and 〈*z*〉 to match the experimental data. The model results, at a reduced bending modulus (*k* = 0.1, solid green line), capture the effect of crosslinker inhibition by T101 treatment on the measured shear modulus (experiments, yellow squares). Both experimental data and model prediction in this case exhibit a noticeable softening dip at ~0.2 Pa. **(D)** Increasing the average coordination number 〈*z*〉, realized in simulations through less random bond removal (*p* denotes probability of bonds being present), leads to higher shear moduli due to greater number of bond springs, and concomitantly reduces the critical strain at which the network abruptly transitions to a stiffer regime. Thus, the position of the transition point in strain can be used to determine the 〈*z*〉 appropriate for the experimental data.

Plasma clots were then prepared between the plates of a rheometer for a subsequent shear assay (see *Methods*). Clot gelation was monitored at low oscillatory strain (<1%) at low frequency of oscillation (1 Hz) in a linear viscoelastic regime. The elastic storage modulus (*G’*) increased over the loss modulus (*G”*) during gelation and reached a saturation value of about ~50 Pa after 45 minutes (Fig. S1). Because of abundant exogenous thrombin, the lag phase typical of clot initiation was not observable, but the rapid lag phase due to fibrin polymerization was followed by a saturation, indicating the formation of a stable clot structure. Interestingly, T101 treatment reduced the initial clot stiffness (at time *t* = 0) but had no impact on the fibrin polymerization rate as indicated by constancy in the slope of the lag phase. Overall, PPP clots treated with T101 demonstrated up to threefold lowering in stiffness in the steady state compared to untreated clots.

Following the initial period of gelation, we recorded the storage modulus *G’* of PPP clots as a function of the applied amplitude of shear stress (data shown as red squares in **Fig.** 2C). The loss modulus *G’’* remained low indicating that the mechanical response is primarily elastic. At low shear stress (< 1 Pa) the clot stiffness stayed constant, but with increasing shear (up to 10 Pa), there was a significant ~3-fold increase in stiffness. In our experiments, we observed that at stresses higher than 70 Pa, *G’* decreased rapidly, indicating irreversible plastic deformations (data not shown). The T101-treated, uncrosslinked PPP clots (data shown as yellow squares) also showed a near constant shear modulus at low shear stress (< 0.1 Pa), and stiffening at high stress (~ 10 Pa). However, at intermediate stresses between 0.1 Pa to 1 Pa, unlike the crosslinked clots, the uncrosslinked clots showed a noticeable decrease in shear modulus by ~10 Pa. For comparison, we also examined the response of fibrin gels formed from 3 mg/mL fibrinogen by the addition of 1 U/mL thrombin and 20 mM CaCl2; and from uncrosslinked fibrin gels formed in the presence of 100 μM T101. As shown in Fig. S2, the response of crosslinked fibrin gels to shear stresses was similar to that of crosslinked PPP clots: strain-independent linear regime at low applied stress, and strain-stiffening at high applied stress. The uncrosslinked fibrin gels showed a response similar to uncrosslinked PPP clots: strain-independent, strain-softening, and strain-stiffening behavior at low, intermediate, and high stress, respectively. The reproducibility of this mechanical behavior between plasma clots and fibrin gels suggests that this is a characteristic elastic response of the fibrin network, independent of other blood proteins.

To explain the highly nonlinear strain-dependent softening and stiffening in crosslinked and uncrosslinked clots, we simulate the response of our model elastic network to shear stress (see *Methods*). There are two important reduced dimensionless parameters in the network model: the mechanical properties of the fibers, namely the ratio between bending and stretching stiffnesses, *κ*/(*μl*_0_^2^), where *l*_0_ is a characteristic fiber length scale given by the distance between branch points; and the connectivity between the fibers in the network, represented by an average coordination number 〈*z*〉. The bending to stretching stiffness ratio *κ*/(*μl*_0_^2^) depends on the ratio of fiber diameter to length, which do not appear explicitly in the model, and is therefore expected to be small for slender fibers. It should be noted that connectivity in fibrin networks is due to the branching of a single fiber into two (or rarely three) fibers, or due to the mutual crossing of two individual fibers leading to entanglement and possible cohesion (*41*). The resulting coordination number (the number of fibers at a branchpoint) is set between 3 and 4 (*41*),(*42*), leading to a dilute, under-coordinated network that has connectivity below the isostatic point (〈*z*〉 = 4 in 2D) for a network of central force springs. The structure of a fibrin fiber is complex (*43*) and includes multiple hierarchical structural levels. To build a minimal model and avoid the structural complexity of fibrin, we treat a fibrin fiber as a single mechanical structure, and estimate the typical bending to stretching ratio *κ*/(*μl*_0_^2^) for a uniform elastic rod (*κ* ~ *Ed*^4^, *μ* ~ *Ed*^2^) using typical, measured values for Young’s modulus (*E*), fiber length (*l*) and diameter (*d*) (*22*). We then allow the bending modulus to be scaled by a free “softening” parameter, 0 < *k* < 1, that is a proxy for the difference of the bending modulus of fibrin, a crosslinked bundle of protofibrils, from the estimated value for a uniform elastic rod. Prior experimental work, that measures the elastic modulus for individual uncrosslinked fibers under bending deformation, suggests that this factor *k* is further reduced when the crosslinking FXIII-A is inhibited (*22*).

The bending stiffness scaling factor k and coordination number 〈*z*〉 were tuned in simulations to match the experimental shear modulus for PPP clots (Fig. 2C). While the shear modulus at small external shear is expected to be dominated by softer, bending modes and thus scales with *k* (*44*), we show that the critical strain for the onset of shear stiffening is relatively insensitive to *k*, instead of depending on 〈*z*〉. This lets us match the flat low shear region, as well as the stiffening transition point, in the experimental *G’* vs. stress curve for PPP clots, by varying *k* and 〈*z*〉, respectively. The fitted value of the bending stiffness factor *k* = 0.4 is smaller than the idealized limit of a uniform elastic rod corresponding to *k* = 1, as expected for a crosslinked bundle of protofilaments (*40*). The coordination number of 3.85 that is found to describe the data well lies between 3 and 4, as expected for fibrous networks. Consistent with our hypothesis that the bending stiffness is reduced for uncrosslinked fibrin, the model captures the response of uncrosslinked PPP clots, for *k* = 0.1. This lowered bending stiffness also led to a softening dip in the shear modulus at intermediate strains, similar to that observed in the experimental data. We explored further the effects of varying network connectivity by simulating the shear response of over- and under-coordinated networks. As shown in Fig. 2D, increasing the average coordination number 〈*z*〉 leads to stiffer networks, while slightly shifting the critical strain associated with the onset of network stiffening to smaller values. This is consistent with the lack of floppy, purely bending modes as the network connectivity increases beyond the isostatic point 〈*z*〉 > 4).

### Shear softening arises from interplay of bending and buckling dominated modes

Since it is experimentally challenging to discern the mechanical state of individual fibers under a given applied shear strain, we use simulations to identify the contributions of different fiber deformation modes to the total elastic energy of the network. These are shown in **Fig. 3** for the bending stiffness factors obtained in Fig. 2C, that corresponds to crosslinked and uncrosslinked fibrin, respectively. We found that both networks comprising fibers with stiffer (*k* = 0.4) and softer (*k* = 0.1) bending moduli exhibit three regimes dominated by fiber bending, buckling, and stretching respectively. Typical simulated, mechanically equilibrated network configurations at representative strain values corresponding to these regimes are shown in Fig. 3A. The different colors represent the type of strain in the corresponding bond, i.e. whether it is stretched (blue), compressed (yellow) or buckled (magenta). Gray (unstrained) bonds can participate in bending modes by changing their mutual angles (not visualized).

**Fig. 3.**
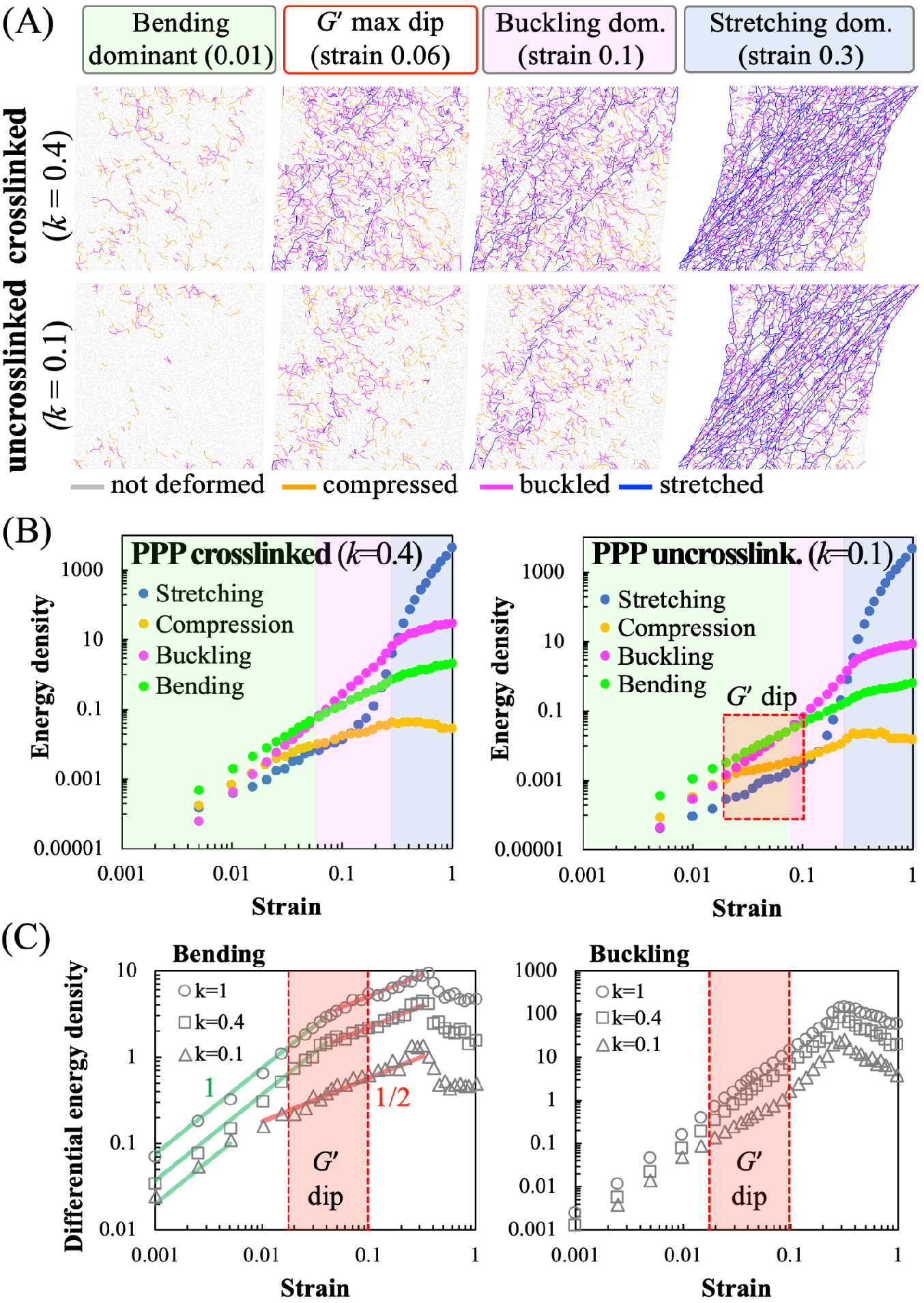
Contribution of different deformation modes to shear response of model elastic networks. **(A)** Simulation snapshots of deformations of crosslinked and uncrosslinked networks at various applied shear strains. Individual bonds in the network are colored according to whether they are stretched, compressed or buckled. The relative occurrence of the different deformation modes depends on both applied strain and fiber crosslinking (corresponds to the value of the bending and buckling stiffness parameter *k*). **(B)** Calculated elastic energy density corresponding to stretching, compression, bending and buckling modes in model simulations for crosslinked (*left*) and uncrosslinked (*right*) cases. The model predicts that both crosslinked (PPP) and uncrosslinked (PPP+T101) clots exhibit three distinct deformation regimes shown by the colored regions, dominated by fiber bending, buckling and stretching, respectively. The red box indicates the regime of strains where the softening dip in the shear modulus is seen in Fig. 2c, and occurs here near the bending-to-buckling transition. This transition occurs at a higher strain for the uncrosslinked/lower *k* network (*right*) resulting in a broader bending-dominated (green) region. **(C)** Differential bending (*left*) and buckling (*right*) energy densities clearly show different power law regimes vs. applied strain. The buckling energy density in the *k* = 0.1 case an additional shallower regime with a lower slope in the region of the softening dip (red box) before approaching with the higher *k* cases. This is consistent with the delayed bending-to-buckling transition seen in **(B)** and fewer buckled bonds in **(A)**.

At very low applied strains (less than 5%), deformations in an under-coordinated network are dominated by bending modes because slender fibers are easier to bend or buckle and incur less elastic energy cost than being stretched or compressed *κ*/(*μl*_0_^2^) ≪ 1). As a result, only a few bonds are seen to be stretched at 1% strain at *k* = 0.4, and the network avoids buckling and compression in bonds at *k* = 0.1. The constant shear modulus in this low strain regime scales with the fiber bending modulus (*34*). At higher applied strains (greater than 10%), individual bonds align along the principal tension direction, resulting in many stretched bonds tilted approximately 45° along the shearing direction. In this regime, the network becomes much stiffer since the stretching energy of individual bonds is high, and a different scaling law of shear modulus with strain results. This is consistent with the well-known rigidity transition from the bending to the stretching-dominated regime under external strain which removes the available bending degrees of freedom (*45*). At very high applied strains, the shear modulus reaches the upper limit of stiffness set by aligned and stretching Hookean springs, and the curves corresponding to different bending moduli converge in the simulations (Figs. 2C and 2D). This limit is not attained in our experiments as the clots start to undergo irreversible plastic deformations at large or oscillatory applied strain (*46*).

Interestingly, for both crosslinked and uncrosslinked networks, there exists an intermediate regime between the low shear, bending-dominated and high shear, stretching-dominated regimes (magenta region in Fig. 3B). In this intermediate regime, the greatest contribution to elastic energy comes from buckled bonds under large compression. In the simulation snapshots in Fig. 3A, this strong influence from buckling is manifested as a relatively high fraction of buckled bonds at intermediate strains (5% to 10%), with the fiber compression oriented approximately transverse to the principal stretching direction. At such low-to-intermediate strain values, the compression of these bonds exceeds the buckling threshold, while the stretching is still relatively small. As seen in Fig. 3B, this intermediate buckling-dominated regime occurs in both the *k* = 0.4 and *k* = 0.1 networks. However, the transition from bending-to-buckling-dominated regimes occurs at slightly higher strains for the *k* = 0.1, compared to the *k* = 0.4 network. This delayed bending-to-buckling transition allows for the intermediate, shear softening region (indicated by red box in **Fig. 3B** (*right)*) corresponding to the dip in the *G’* curves, seen only for *k* = 0.1.

To better understand the origin of this shear softening (seen only at lower *k*) and to identify how the different energy contributions scale with increasing shear strain, we plot the differential energy density against strain for different *k* values in Fig. 3C. The differential bending energy trends show a transition from a steeper (~1) to a shallower slope (~1/2) power law regime, with the transition strain being higher for higher values of *k*. This shift in transition strain indicates that the bending energy grows slower with strain, as buckling, and then stretching, modes take over. Similarly, for *k* = 0.1, a shallow-slope regime is seen in the differential buckling energy. This correlates with the later onset of the buckling-dominated regime for *k* = 0.1 seen in Fig. 3B, when compared to cases with higher values of *k*. Taken together, these suggest that the softening dip in shear modulus is a result of buckling of initially compressed bonds as the strain increases from very low values. For *k* = 0.1, this dip is fully expressed, since the number of stretched bonds is still very small at this strain regime, while the effect is lost at higher *k* when a significantly larger number of bonds gets stretched. Thus, by inhibiting fibrin crosslinking, our experiments offer a direct confirmation of this intermediate shear softening effect, which was previously noted only in simulated elastic networks with buckling (*47*),(*48*),(*49*).

### Fiber crosslinking promotes efficient force transmission in platelet-contracted networks

Having examined the behavior of passive fibrin networks, we next sought to examine the differences in the interaction of platelets with crosslinked and uncrosslinked fibrin networks. The differences in platelet morphology and fiber distribution around the platelets are highlighted in Fig. 4A. As shown in the inset for the PRP clot, the fibers are more uniformly oriented around the platelet cluster, which then extend filopodia along these fibers, resulting in a more isotropic configuration. In contrast, the T101-treated (i.e., uncrosslinked) PRP clot exhibits irregular distribution of fibers around the platelet aggregate, lacks filopodia, and has more anisotropic morphology. It is well-known that platelets exert contractile stress on vicinal fibrin fibers shortly after initiation of the clot which reaches a steady state as the clot is stabilized (*50*). Thus, while the local mechanical deformations of the fibrin network cannot be quantified as yet, the organization of fibers and filopodia are expected to be correlated with local strains.

**Fig. 4.**
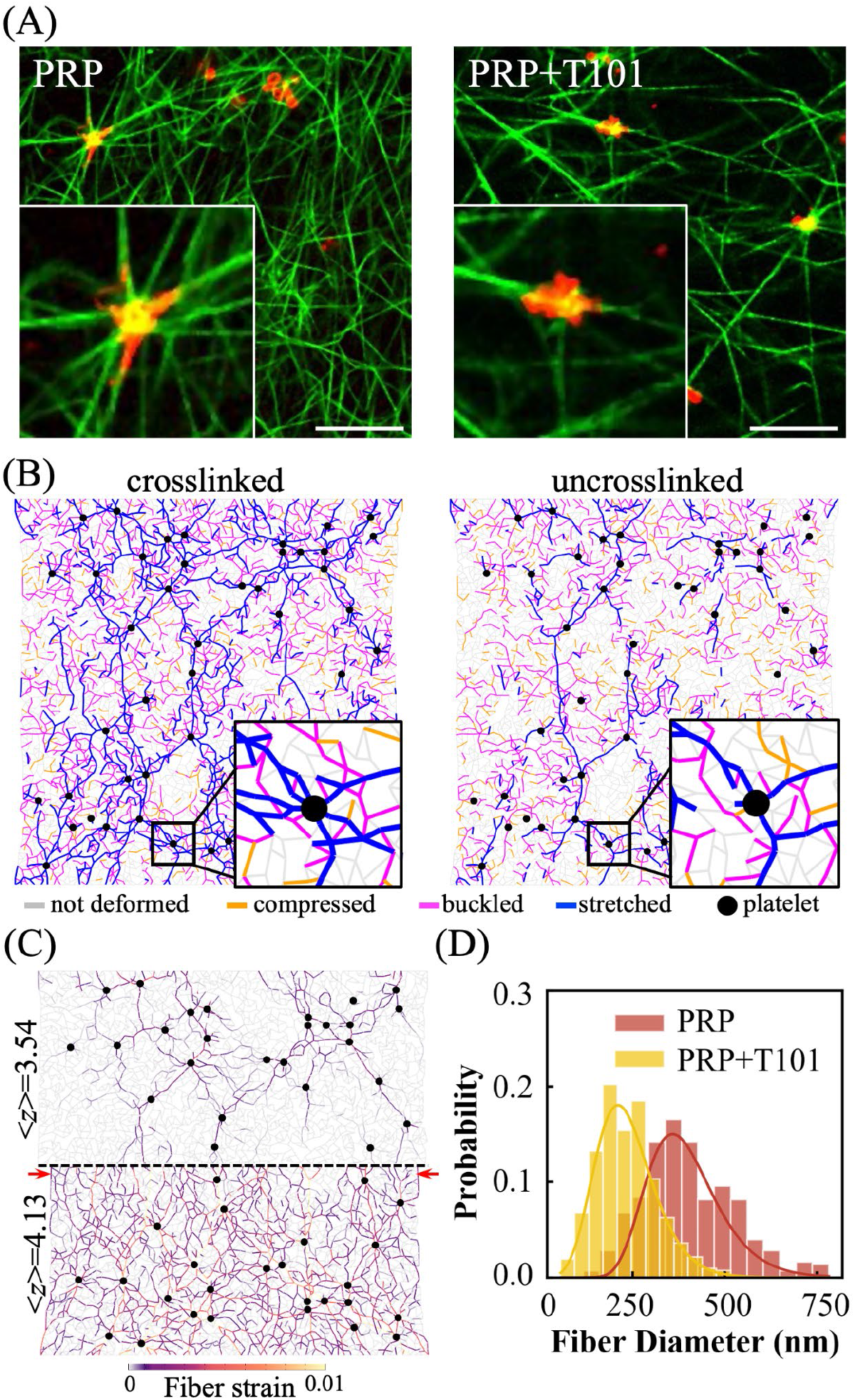
Fibrin crosslinking is critical to platelet-fibrin interactions and to the formation of force chains. **(A)** Fluorescence microscopy images of platelet-rich plasma (PRP) clots with crosslinked fibrin (*left*) and with T101 inhibition of crosslinking (*right*). Insets clearly show that fibrin (green) distribution is more uniform around platelets (red) in the crosslinked clots, with the platelets also showing a more isotropic morphology with pronounced filopodia in these cases. The empirical observations are consistent with model prediction of higher densification for more stretched crosslinked fibers. Scale bar: 100 μm **(B)** Model network simulations show the heterogeneous distribution of forces around actively contractile nodes representing platelets (black). The stretched fibers (blue) form “force chains’ ‘ that propagate from one platelet to another and are sparser and more localized in the uncrosslinked network (*right*). **(C)** Force chains are significantly denser and longer-ranged in over-coordinated 〈*z*〉 > 4) than under-coordinated networks 〈*z*〉 < 4). In both cases of lower bending stiffness (*k*) and lower 〈*z*〉, there are many energetically favorable bending modes available, which lead to reduced fiber stretching. This in turn will lead to reduced force propagation to the network boundary, predicting an anomalous effect of less macroscopic contraction in softer networks (where the retraction difference is shown by red arrows). (**(D)** Quantification of apparent fiber diameter around platelet clusters (*n* = 4 donors; 70 fibers per sample). Thus, inhibition of crosslinking by T101 treatment of PRP clots leads to thinner fibers around platelets. This suggests less bundling of fibers and weaker platelet-fibrin interactions.

To connect the fiber distribution and platelet morphology to local deformations, we simulated the response of the elastic network to active contraction, by introducing actively contractile platelets into the passive fibrin network (see *Methods* for details). Uncrosslinked and crosslinked active networks were modeled by modifying the bending-to-stretching stiffness ratio, represented by the stiffness factor, *k*. Our previous experimental estimate of about 140 platelet aggregates/mm^2^ in a cross-section of contracted clots (*5*) was used to seed the network with ~70 active nodes in the simulation box size which corresponds to an area of 0.5 mm^2^. The contractile forces exerted by the platelet aggregates were obtained from published estimates from micropost measurements as ~30 nN (*51*). The contractile prestress exerted by the platelets on the fibers manifests as an increase in the stiffness of PRP clots compared to PPP clots. The gelation kinetics showed that the presence of platelets increased the stiffness of clots even at the onset of gelation (Fig. S1). In fully formed clots, platelets increased the shear modulus of the plasma clots by threefold, while a twofold increase due to platelets was also seen in uncrosslinked clots.

In our active network model for PRP clots, the platelet aggregates are modeled as isotropic force dipoles that continuously generate contractile force. This is realized in the simulations by reducing the reference length of bonds attached to the active nodes, representing platelet aggregates. The equilibrium configuration is then achieved when the forces exerted by the platelets are balanced by equal and opposite elastic restoring forces from the bonds representing fibrin fibers. The simulation results showed that the platelet contraction does not increase the strain uniformly in all fibers but only in some, which together form an interconnected subnetwork of stretched fibers highlighted in blue (**Fig.** 4B). We observed a denser subnetwork for fibers with higher bending rigidity, i.e., corresponding to crosslinked fibers. In case of uncrosslinked fibers with lower bending rigidity, the high-strain subnetwork is sparse, and appears as a few long and connected chains. This difference is again because more bonds can easily rotate to relax their tension at lower values of *k*. The subnetworks of stretched fibers or “force chains” originate from and extend to distant platelet aggregates, and thus serve as conduits for the transmission of forces exerted by the platelets. Consequently, the crosslinked PRP clots which are made of fibrin fibers with high bending rigidity and denser force chains, are expected to contract more than the uncrosslinked clots (*28*).

Importantly, and for the same underlying reason as networks with stiffer bending (high *k*), networks with higher coordination number also exhibit denser force chains (Fig. 4C). The availability of low energy bending modes in under-coordinated networks (lower 〈*z*〉), causes the stress generated by platelet aggregates to be localized only along a few force chains. In denser networks at larger average coordination number 〈*z*〉, stresses propagate more uniformly throughout the network (more fibers under strain), and as a result, the range of force transmission and bulk network contraction is expected to be larger. The difference in the range of stress propagation is qualitatively seen at the midline of the network depicted by the dashed black line in **Fig. 4C** (only mirror halves of networks are shown). Clearly, many more force chains reach the midline for the higher 〈*z*〉 case. We chose 〈*z*〉 = 4.13 in the simulation for representational purposes to show this marked contrast, although we do not expect fibrin networks to exhibit an average coordination number greater than 4. Although it is difficult to directly quantify from experimental images how the coordination number of fibrin networks in clots changes upon T101 treatment, it is plausible that crosslinker inhibition may reduce the number of branch points along with the bending stiffness at these branchpoints. Thus, our model predicts that crosslinker inhibition will lead to fewer force chains, less efficient transmission of forces in the network, and less overall retraction of the clot. This behavior can be detrimental to the strength and stability of fibrin networks, and can lead to clot failure.

To elucidate the fiber redistribution around platelets revealed by the experimental micrographs (insets in Fig. 4A), we focus on the force chains around active nodes shown for the simulated networks as insets in Fig. 4B. The fibers directed radially outwards from the active nodes transmit the highest tensile strain (blue), while those along the azimuthal direction tend to be compressed, and then buckled (purple). The stretched fibers are more numerous for the higher *k*-network, as expected, due to larger bending and buckling resistive forces from other fibers. In contrast, simulations show fewer force chains for the network with lower *k*, which travel along fewer directions. To relate this observation to experiments, we quantified in **Fig. 4D** the apparent diameter of fibers around the platelet aggregate using images such as in Fig. 4A. These fibers close to platelets were found to be thicker in the crosslinked case, with mean and standard deviation values of 402 nm ± 126 nm for PRP clots and 236 nm ± 81.5 nm for PRP clots treated with T101 (*n* = 4 donors with about 70 fibers per donor; *P* < 0.001, two-tailed t-test). Our model predictions for the force distribution around platelets (Fig. 4B) suggests a plausible explanation for this observation. The more numerous force chains that develop between two nearby platelet aggregates in the stiffer crosslinked networks, can lead to the alignment and bundling of fibers between them, which may be further stabilized by FXIII-A-mediated lateral crosslinking. Contractile cells (fibroblasts) in fibrin gels have been shown to form such densified and aligned bands of fibers around them, through which they mechanically interact (*17*). For uncrosslinked fiber networks, our model suggests that there are fewer force chains between platelet aggregates, which will therefore lead to weaker bundling of the fibers and lower effective fiber diameter, as observed. This hypothesis is further supported by the observation that the diameter of the fibers far from the platelets is comparable between crosslinked and uncrosslinked clots, demonstrating that platelet activity was responsible for fiber bundling (*52*),(*53*). Altogether, the results in **Fig. 4** show that inhibition of fibrin crosslinking strongly impacts the active remodeling of the surrounding fibrin network by platelets.

### Active stiffening is dependent on magnitude of shear stress and extent of crosslinking

Next, we examined the mechanical response of crosslinked and uncrosslinked PRP clots to shear stress (**Fig. 5A**). PRP clots were prepared between the plates of a rheometer, allowed to gel, and then subjected to shear. To simulate this process, we first allowed the model networks to contract under active forces, as described in the previous section, but now with nodes on the network boundary held fixed. Subsequently, these clots were subjected to shear strain as described previously for PPP clots, but now with randomly placed “active nodes” representing platelet aggregates that actively pull on the fiber network. The PRP network simulation results capture the experimental results with the same coordination number as the passive networks 〈*z*〉 = 3.84), but with a modest increase in the bending stiffness parameter, *k*. This may indicate that fibers become stiffer to bend in the presence of platelets, possibly through platelet-induced crosslinking. The comparable estimates of model parameters in the passive and active networks from the low and moderate shear regimes, suggest that platelets at this density primarily modify the mechanical prestress in the network, and not its connectivity 〈*z*〉 or fiber structure. Overall, the response of PRP clots is similar to PPP clots, and shows a stiffening transition at large strains. However, unlike PPP clots, platelets not only stiffen the network but also cause a softening dip of different magnitudes in both crosslinked and uncrosslinked networks (**Fig. 5A**). The softening regime in PRP clots occurs at about the same strains as in PPP clots, and is also more pronounced in the uncrosslinked networks.

**Fig. 5.**
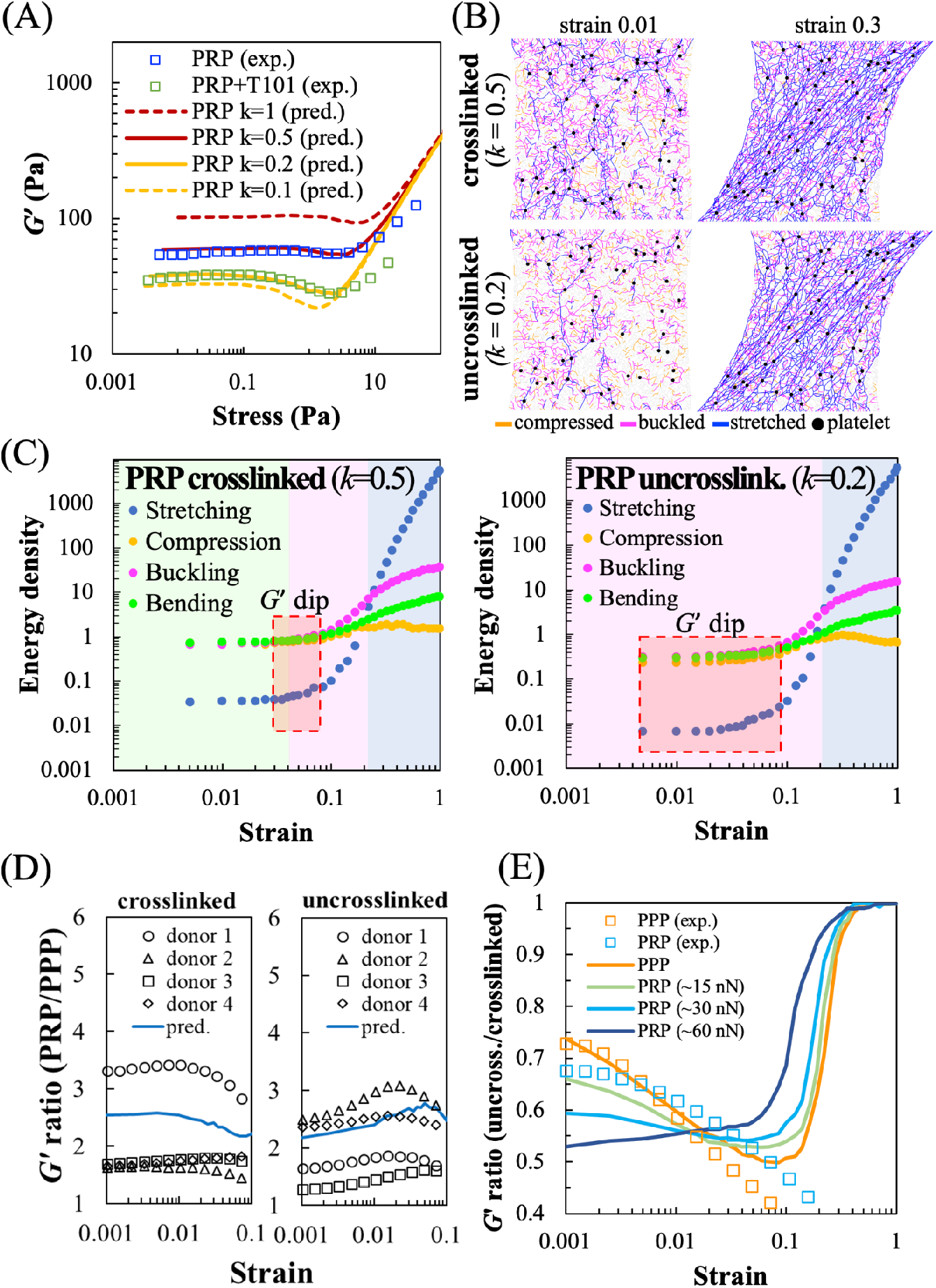
Shear stiffness of active networks representing platelet rich plasma (PRP) clots. **(A)** Shear modulus (*G’*) vs. applied stress for PRP clots (squares), with and without T101, and for simulated networks (lines) at <z>=3.84, ~30 nN initial platelet contractile force, and different values of the reduced fiber bending stiffness parameter, *k*. Uncrosslinked PRP clots demonstrate a pronounced softening dip, unlike PPP. **(B)** Simulation snapshots of network deformation vs. strain, with individual bonds colored by strain. The number of stretched bonds is high due to platelet-generated prestress and increases further with applied shear strain. **(C)** Calculated elastic energy density, corresponding to the different deformation modes, as a function of applied strain in simulations. The colored regions indicate the deformation mode that is the largest contributor to the total energy, while the red dashed box indicates the region where the softening dip is observed in the shear modulus in **(A)**. **(D)** Ratio of shear moduli of PRP to PPP clots at different applied shear strains, shown for four different donors (data points) and corresponding simulations (line), for both crosslinked (*left*) and uncrosslinked (*right*) cases. Thus, platelets stiffen the network by different amounts depending on applied shear. The stiffening factor decreases gently with strain for crosslinked networks while for uncrosslinked networks, it increases with strain up to a maximum at 6%. **(E)** Ratio of shear moduli of uncrosslinked to crosslinked networks in simulations (lines) for different values of platelet contractility (initial force values shown), and correspondingly, ratio of measured shear moduli of T101-treated to untreated clots (squares). Strongly contractile platelets (~60 nN) amplify the difference and show different behavior vs. strain.

**Fig. 6.**
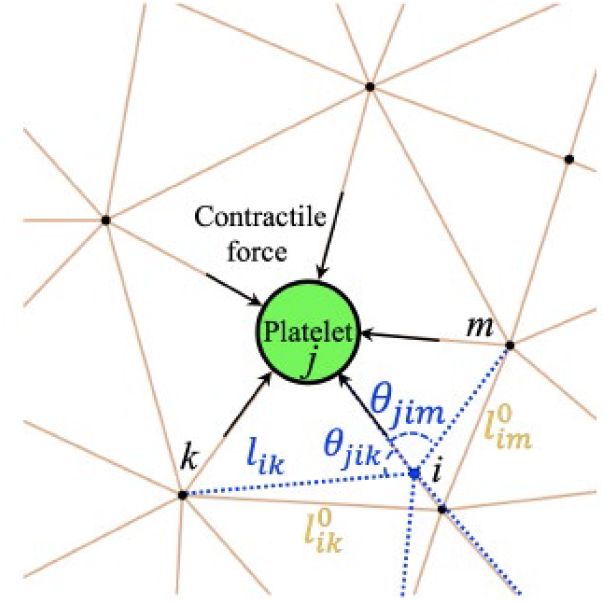
Geometry of the elastic network model. The model comprises uniform spring-like bonds connected at nodes of different coordination. Some nodes are “active” and represent aggregates of platelets generating contractile forces on connected fibers. Under the action of active nodes and applied strain, the initial rest configuration (yellow bonds) deforms to a new stressed configuration (blue bonds) by displacing connected nodes. This deformation is associated with changing fiber’s end-to-end lengths (stretching/compression) and changing the angles between bonds (bending). Both these deformation modes result in an elastic restoring force on the concerned node. The calculated forces lead to the overdamped dynamics of the node towards the mechanical equilibrium state where all forces are balanced.

To gain qualitative insight into the different regimes of the shear modulus vs shear stress curves in **Fig. 5A**, we show representative snapshots of the simulated active networks at low and high strains in **Fig. 5B**. Individual network bonds are colored to show their deformation state, i.e. whether it is stretched (blue), compressed (yellow) or buckled i.e. compressed beyond the critical strain (magenta). Unlike similar snapshots for networks without platelet contraction shown previously in **Fig. 3A**, these networks have considerable stretched and buckled bonds even at the very low strain value of 1%, due to the prestress induced by platelets. The number of stretched bonds increases rapidly with applied stress, because the prestress pulls out floppy bending modes. The corresponding bending-to-stretching stiffening transition happens at lower applied strains in the presence of platelets both in uncrosslinked (**Fig. S4A**) and crosslinked networks (**Fig. S4B**), as a result of this. The force chains propagating from the platelets intensify with applied strain (**Fig. 5B**), but they remain more localized to the paths connecting platelets in uncrosslinked networks. The effects of both crosslinking and platelet prestress are eliminated in the stretching dominated regime at larger strains, e.g. at 30%. Similarly to **Fig. 3B** for networks without platelets, we now consider in **Fig. 5C** the relative contributions to elastic energy density of the different deformation modes. We observe two distinct differences between active networks with platelets and passive networks without platelets (**Fig. 3C**). First, the platelet-induced prestress creates an initial plateau of energy for all deformation modes at low applied strain. In this initial regime, the effect of the platelet prestress dominates that of the applied stress. Second, unlike passive networks which have predominantly bending modes at low strains, the platelets induce compression and buckling with energies comparable to that of bending **(Fig. 5C)**. While the crosslinked network at larger *k (left*) still has a bending-dominated regime (**Fig. 5C**), this regime does not appear in the uncrosslinked, lower *k* network (*right*). The absence of this regime is again due to platelets, which are more effective at pulling out floppy modes in uncrosslinked networks that have smaller bending forces.

While it is expected that platelets will stiffen the fibrin network by introducing prestress, we next examine the magnitude of this stiffening. The stiffening effect of platelets on clots is seen in terms of the ratio of the shear modulus *G’* for PRP to PPP, shown separately for uncrosslinked and crosslinked networks (**Fig. 5D**). Both the measurements for individual donors (discrete markers) and the model calculations (solid line) show a significant increase (about twofold to threefold) in shear modulus produced by platelets. Interestingly, the stiffening is shear-dependent for both crosslinked and uncrosslinked networks. In crosslinked networks, the stiffening due to platelets is most appreciable at low strains, and decreases weakly with strain. On the other hand, this stiffening in uncrosslinked networks exhibits a noticeable maximum at intermediate shear. This is consistent with the model prediction that platelets stiffen most at strains where the maximum softening dip in *G’* occurs for PPP (~6%). By pulling on the surrounding fibers, platelets stiffen the network through a reduction in the number of floppy, low-energy bending modes. In the high strain regime, however, the applied external force becomes much larger than the force generated by platelets. Thus, at high strain PPP and PRP clots should have the same value (i.e., *G’* ratio trending toward 1). To summarize, the results in **Fig. 5D** indicate that platelet-fibrin interactions depend on both the amount of applied stress and the fiber bending stiffness associated with fibrin crosslinking. The larger stiffening effect of platelet-generated prestress in uncrosslinked networks suggests that prestress may serve as a compensating mechanism for restoring clot stiffness in the absence of crosslinking. The platelets are most effective and provide sufficient clot stiffening at small and moderate strains (up to 10% strain), which may be physiologically relevant at early stages of clot formation.

To examine the effect of prestress due to platelets on networks with and without crosslinking, we plot the ratio of shear moduli of uncrosslinked to crosslinked clots predicted from simulations, at different values of platelet contractility (**Fig. 5E**). The corresponding shear moduli for simulated networks at higher and lower bending stiffness, are shown in **Fig. S4D and S4E**. Unlike PRP clots, the PPP data (orange squares) show closer agreement with simulations (orange line) at low-to-intermediate strain (<10%). This suggests that platelets induce strain-dependent changes to network mechanics, which are not fully captured by our simple model which does not strictly control platelet contractility. In the simulation curves in **Fig. 5E**, all ratios start less than 1 in the bending-dominated low strain regime, and approach 1 at much higher strain values where stretching dominates and the differences in *k* are not relevant. In the intermediate regime, the shear moduli ratio is sensitive to the contractile forces generated by platelets. For platelets with weaker contractile forces (half the estimated initial value, *~*15 nN (*51*)), the pronounced dip in the *G’* vs. strain curves for uncrosslinked networks leads to an initial decrease and subsequent increase in this ratio. However, stronger platelet contraction (double the estimated initial value, ~ 60 nN) shows greater softening for the uncrosslinked networks before external shear is applied. Additionally, platelets with higher contractility express a softening dip in the *G’-*vs-strain curves for both crosslinked (*k* = 0.5) and uncrosslinked (*k* = 0.2) networks (see **Fig. S4D** and **S4E**). This behavior is different from networks with lower platelet contractility which show a pronounced softening dip only at lower *k* values. The stronger platelets soften the network more when crosslinking is inhibited at small strains than at intermediate strains. Taken together, the simulation curves in **Fig. 5E** show that the manner in which platelet contractility influences modulus (specifically the *G’* ratio for uncrosslinked vs. crosslinked fibers) is dependent on the magnitude of strain. This may arise due to the adaptive dependence of platelet contractile force on network stiffness (*54*), through which platelets possibly exert lower initial force, e.g. 15 nN, at low strains (below 1%), and exert higher force at intermediate strains (1% to 10%). Such mechano-adaptation is typical in contractile cells that exert greater traction force when the extracellular environment is stiffer. Additionally, we note that the simulation curves predict a crossover in the ratio of shear moduli where these values become identical for networks at different contractility (~2%). This crossover change in slope—seen in both simulation and experimentation—suggests that there is a strong interaction effect between platelet force and magnitude of strain.

## DISCUSSION

In this work, we combined modeling and experiments to characterize the macroscopic mechanical responses of crosslinked and uncrosslinked plasma clots to shear strain in the presence and absence of active prestress induced by platelets. To understand the micromechanical origins of these responses, we resolved the contribution of individual fiber deformations to the overall mechanical behavior using a minimal elastic network model parameterized from experiments. Our results show that the network response at various shear strains is dominated by different deformation modes of individual fibers, and depends on fiber crosslinking, network connectivity, and platelet contraction-induced network prestress. Our model predicts that the experimentally observed unusual shear softening transition occurs due to the propensity of uncrosslinked fibers to buckle and bend rather than to be stretched or compressed. Our model can thus capture the effect of a biochemical perturbation-induced change in fiber structure through a single coarse-grained parameter (*k*) corresponding to fiber buckling and bending, thereby linking molecular scale structure to a continuum mechanical property (*42*).

While the rheology of fibrin gels are well characterized (*55*), understanding the physical mechanisms governing behavior of plasma clots, particularly the effects of fiber crosslinking and platelet-induced contraction, remains elusive. Fibrin gels stiffen under increasing shear, although the extent and nature of stiffening vary depending on the polymerization conditions. Consistent with previous reports, we show that fibrin gels and PPP clots transition from linear response characterized by a constant shear modulus to shear-dependence at ~10% strain or at a stress of ~1 Pa (*56*),(*57*)]. This strain-stiffening phenomenon is well-established in other biopolymer networks such as actin and collagen (*10*) and in fibrous networks in general (*58*). These typically comprise semiflexible polymers that show nonlinear force-extension curves with stiffening under tension which pulls out thermal fluctuation modes, and softening under compression. Semiflexible polymer networks are expected to stiffen according to a specific power law with shear stress, *G’* ~ *σ*^3/2^ (*59*), as has been demonstrated in the case of crosslinked actin gels (*60*). Strain-stiffening may also arise as a collective effect in athermal fiber networks, due to the purely geometric effect of strain-induced fiber alignment (*61*), as well as the strain-induced transition from a bending-dominated to stretching-dominated response in under-coordinated networks (*45*). To investigate the nonlinear strain-response regimes of plasma clots, we developed and compared experiments with a general, enthalpic model of fibrin networks wherein the network mechanics is governed by the bending, buckling, and stretching modes of the constituent fibrin fibers (*37*). At higher strains where clot rupture occurs, the irreversible dissociation of bonds between fibrin monomers might become more important, but we ignore this effect in our model since we aim to capture experiments at low to intermediate shear where network response remains reversible and elastic (*62*).

The agreement between model predictions and experimental measurements of shear modulus at different applied strains demonstrates that the bending and buckling modes of fibers are important for describing the shear response of fibrin networks. Individual fibrin fibers buckle as much as one-third their unstrained length under compression, and they stretch nearly two times their original length under tension (*63*),(*23*). Under compression, clots may expel fluid leading to poroelastic effects that are not considered in the present model (*64*). We focus here on the elastic response of the clots and not plastic or possible viscoelastic effects which are undeniably important for understanding the full regime of clot dynamic behavior. Elastic energy calculations show that the mechanics of the network are governed by bending, buckling, and stretching with the dominant mode strongly dependent on applied strain. The rate of growth of constituting elastic energies with strain exhibit different regimes characterized by different power laws that depend on the bending/buckling stiffness parameter, *k*. With increase in applied strain, two distinct transitions in predominant modes of deformation occur, namely bending-to-buckling, and buckling-to-stretching. The latter is manifested as an abrupt increase in *G’* independent of crosslinking, and this transition is well-documented as the strain-stiffening response of biopolymer networks (*13*). Although softening of fiber networks has been reported in simulations (*47*),(*48*) in this work we provide the first experimental proof of this effect in crosslinking-inhibited plasma clots and also show that this arises from the bending-to-buckling transition which occurs at low strains. This effect may be a broader phenomenon: recent rheological measurements of clots from rats (*65*) reported a significant decrease in shear modulus with increasing shear deformation, although the mechanistic or mechanical basis of such softening response was not investigated. Here, by comparing the mechanics of uncrosslinked and crosslinked PPP networks, we show that the atypical softening-stiffening behavior of uncrosslinked fibers originates from a bending to buckling-dominated response, which softens a fraction of the fibers originally under compression.

We explored heterogeneous force transmission in the elastic network subjected to externally applied shear or due to internal platelet contraction. Our simulations show that the fibers transverse to the direction of external load are under compressive stresses and they eventually bend and buckle; while fibers longitudinal to the direction of external load are under tension and get stretched. This orientational anisotropy gives rise to spatial heterogeneity in microscale stress distributions which govern their overall behavior including non-linear stiffness and tendency to rupture (*66*),(*67*). The local heterogeneity in strain distribution also manifests as disordered patterns of force transmission through tensile force chains (*17*),(*18*),(*68*). Force chains are believed to be the conduits for long range force transmission between contracting cells in fibrous materials (*69*). Unlike previous models that considered only fiber buckling in the analysis of force chains that develop between contractile cells in fiber networks (*24*),(*70*), here we include both bending and buckling of linear elastic fibers in our modeling, and show that bending screens out long-range propagation of strains in the network. We thus find that the decrease in bending stiffness of fibers results in fewer force chains that extend through the network. Likewise, we observed more numerous and longer force chains connecting platelets in higher bending stiffness networks corresponding to crosslinked fibrin. We connect this to the experimental observation of thicker aligned bundles of fibers around platelets in a crosslinked fibrin network, and a more isotropic and well-spread platelet shape. Lastly, our analysis suggests that a lack of sufficient number of force chains may result in an inability to transmit the contractile forces generated by platelets in uncrosslinked PRP clots, which provides an explanation for our previous experimental observation that T101-treated PRP clots failed to generate measurable contractile force (*28*). Thus, we predict anomalous elastic behavior in contracting networks: stiffer fibers result in greater contraction under internal active stresses, which is counter-intuitive when compared with a solid under external compression.

Importantly, we have investigated the response of the fibrin networks to shear stresses which are well within physiological and pathophysiological ranges: the wall shear stresses in blood vessels range widely between 0.1 Pa (in veins) to 10 Pa (in stenosed arteries and arterioles) (*30*); indirect estimates of stresses at various regions of blood clots are reported to vary widely between 0.1 to >100 Pa (*71*),(*72*); and the shear stresses experienced by thrombi may increase by several fold during stenosis (*73*) or due to local variations in clot porosity (*74*). Therefore, the increasing stiffness beyond 2 Pa may serve to mitigate spontaneous mechanical damage to the clots in regions of high shear stress. Strain stiffening of fibrin networks is a remarkable phenomenon that may have evolved to maintain the integrity of blood clots and fibrin sealants under large deformation such as when exposed to shear stresses due to blood flow.

The implications of our findings at stresses lower than 2 Pa might be particularly important at earlier stages of clot formation. Stresses below 2 Pa arise not only in slower blood flows but also in the remodeling of fibrin networks during cell contraction (*75*). At these low shear stresses, we noticed qualitatively different behaviors of crosslinked and uncrosslinked PPP clots (Fig. 2), and the maximum shear stiffening effect of platelets on uncrosslinked, but not on crosslinked, PRP clots (**Fig. 5**). Of consideration is the timing of FXIII-A-mediated crosslinking in the formation of mature, fully contracted clots. Since the formation of laterally aggregated protofibrils always precedes their crosslinking, weakly compacted and uncrosslinked bundles of protofibrils are simultaneously remodeled by FXIII-A and by platelets (*76*),(*77*). The pronounced softening of uncrosslinked fibers observed in our experiments suggest that strain-dependent stress propagation through these fibers will impact the final clot structure. Our results also suggest that softer fibers easily bend and buckle under compression, which makes them acquiescent to easy rearrangement but not to long-range force transmission, indicating a location-dependent remodeling in evolving clots. Hence, the timing and extent of FXIII-A-mediated crosslinking dictates clot structure not only through the direct modification of fiber thickness and branching but also by altering the trajectory of the stiffness landscape.

Overall, we have described a general elastic network model that incorporates platelet activity, and can accurately capture the shear response of PPP and PRP clots. The model accounts for different fiber deformation modes (stretching, compression, bending, and buckling), the extent of fibrin crosslinking, and the contractility of platelets. We have also described a potentially overlooked softening regime in fibrin networks at intermediate shear, before the well-known onset of strain stiffening. The softening is shown to arise due to reduction in bending stiffness of uncrosslinked fibers and a delayed bending-to-buckling transition. This softer regime may arise in early stages of clot formation, allowing for greater restructuring by platelets. Lastly, we show that crosslinking also modulates the distribution of prestress imposed on the network by platelet contraction, and changes the deformation modes of the network to applied shear. Our work provides biophysical insights into how mechanical cues from the fibrin network, external stress and platelet contraction work together to modulate macroscopic clot stiffness, and provokes the possibility of adaptive mechanical regulation in the clotting process.

## MATERIALS AND METHODS

### Experimental methods

#### Isolation of platelet rich plasma (PRP) and platelet poor plasma (PPP)

Blood was obtained by phlebotomy from healthy volunteers between 20 to 30 years of age who did not have any chronic conditions or medications known to alter platelet function (San José State University IRB protocol F16134). The blood was drawn in vacutainer tubes containing 3.2% buffered sodium citrate (BD Biosciences, San Jose, CA, USA). The blood was centrifuged at 250 RCF (relative centrifugal force) and 7 rad/s^2^ acceleration for 20 min (5810 R, Eppendorf, Hamburg, Germany). The platelet rich plasma (PRP) supernatant was separated from the red blood cell (RBC) sediment. To obtain platelet poor plasma (PPP), 10 μM PGI_2_ (Sigma) was added to PRP, and was further centrifuged at 600 RCF and 9 rad/s^2^ acceleration with brakes for 15 min. The supernatant PPP was separated from the platelet pellet. For rheometry experiments, the PRP and PPP were used within 4 and 6 h of isolation, respectively.

#### Shear rheometry of PRP and PPP clots

Shear experiments were executed on 600 μL of PRP or PPP mixed with 1 U/mL thrombin (Enzyme Research Laboratories, South Bend, IN, USA) and 20 mM CaCl2 (Millipore Sigma, St. Louis, MO, USA) subjected to shear strain at 22 °C in rheometer in two stages (MCR 302, Anton Paar, Graz, Austria). First, a small amplitude oscillatory test at low strain (0.5%) and 1 Hz was applied for 45 min. Second, after clot gelation, shear strain was progressively increased from 0.001% to 250% at a constant frequency of 1 Hz. The shear stress and shear modulus were recorded. To prevent evaporation, a thin immiscible oil layer (Vapor-Lock liquid vapor barrier, QIAGEN, Valencia, CA, USA) was applied along the rim of the clot, and the set-up was covered with a humidifying chamber. For experiments involving inhibition of factor XIII (FXIII-A)-mediated crosslinking,100 μM of T101 (Zedira GmbH, Darmstadt, Germany) was added to PRP or PPP before performing the experiments.

#### Clot microstructure

To prepare PRP clots for visualization, 100 μL of PRP was incubated with 1 μg/mL AF647-conjugated CD42b rabbit anti-human antibody (BioLegend, San Diego, CA, USA) to label the platelets, and 1% AF488-conjugated fibrinogen (BioLegend, San Diego, CA, USA) to label fibrinogen for 10 min on a rocker with gentle rocking frequency of approximately 2 Hz (Vari-mix Platform Rocker, Thermo Fisher Scientific, Waltham, MA, USA). After incubation, clots were prepared by adding 20 mM CaCl2 and 0.2 U/mL thrombin. Immediately, 50 μL of the mixture was added on ethanol-cleaned 1 mm microscope slides (Thermo Fisher Scientific, Waltham, MA, USA). PPP clots were prepared in a similar fashion but without the platelet antibody. The clots were imaged by confocal microscopy with an Apochromat 63X oil objective with a vertical stack interval of 1 μm (Zeiss LSM 700). The images were analyzed using Imaris v9.5 (Oxford Instruments, MA), and NIH ImageJ (*78*).

### Computational model

#### Network model setup

The initial network was generated with Delaunay triangulation in a two-dimensional square domain using the software tool Gmsh (*79*). The triangulated network had an average spacing *l*_0_ = 〈*<u>l_ij_</u>*〉 between neighboring nodes *i* and *j*, and consisted of about 5000 nodes arranged in a box of size 70 × 70 node spacings. The initial network configuration without platelets is assumed to be stress free. Each bond in the network is then at its rest length, *<u>l_ij_</u>*, and each junction at its rest angle, *<u>θ</u>_ijk_* initially (**Fig. 1A**). The average node spacing was set to correspond to the approximate average branch length, 10 μm, typically seen in experimental images. Image analysis based on fluorescence microscopy showed no significant difference between fiber lengths for clots with and without the T101 crosslinking inhibitor (**Fig. S3**). The coordination number was varied by randomly removing a fraction of bonds in the initially fully coordinated network. To prevent dangling bonds, the coordination number at each connecting node was not allowed to be smaller than 3. The probability of a bond being present, *p*, determines the average network coordination number 〈*z*〉. To increase the network irregularity, the position of each node *x_i_* was additionally perturbed by a small random displacement, *δx_i_* = 0.1 *η l*_0_, where *η* is a uniformly distributed random number from the interval [−1, 1]. The active nodes representing platelets were randomly placed throughout the mesh to reach a typical platelet seeding density. To prevent large local deformations in the network caused by locally biased density or a lack of connected fibers at the boundary, active nodes were not allowed to be at the boundary and two platelets were not allowed to share the same bond.

#### Network model dynamics

The dynamics of the simulated network mimics the inertia-free, and viscous dominated clot dynamics. Thus the position of each node in the network was updated following equations for noise-free and overdamped dynamics designed to find the mechanical equilibrium state,

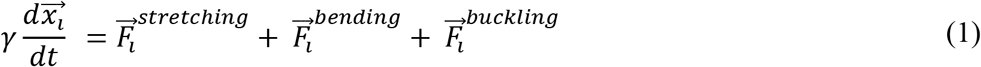

where *γ* is the drag coefficient arising from friction due to the surrounding viscous medium, and the total force acting on the *i^th^* node includes contributions from bond stretching, bonds bending i.e. changing their relative orientation at junctions, and bond buckling forces, given by

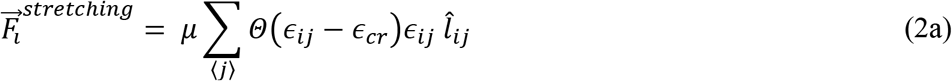

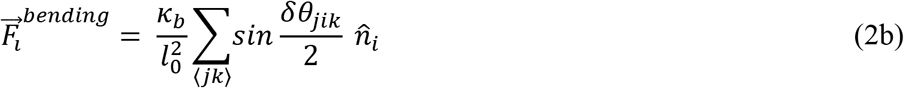

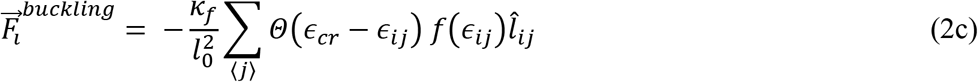

Here, *μ* is the stretching modulus of individual bonds, *Kb* is the bending modulus at junctions between two bonds which penalizes change in the bond angle and can include contributions both from the entanglement between neighboring fibers, as well as the bending of individual fibers. 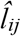 represents the unit bond vector connecting the *i^th^* to the *j^th^* node, and the strain in this bond is given by *ϵ_ij_* = (*l_ij_* / *<u>l_ij_</u>* – 1), where *l_ij_* is the actual bond length, and *<u>l_ij_</u>* the target rest length. The direction of the bending force is along 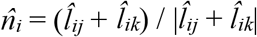. The buckling force arises when the critical compressive strain is exceeded, *i.e*. *ϵ* < *ϵ_cr_*, and opposes the compression. It scales with the bending modulus of an individual fiber, *κ_f_*, and is a nonlinear function of compressive strain,*f*(*ϵ_ij_*) (*80*). The Heaviside function Θ takes the value Θ = 1 when its argument is positive, and Θ = 0 in the other cases, and therefore, determines if the fiber is stretched, compressed or buckled. In our model, we do not calculate the bent shape of the fibers, but instead model the effect of buckling as a force that resists compression, with an effective modulus that is much smaller than the stretching modulus, *μ*. Further, while the bending stiffness of individual fibers *κ_f_* can be different in principle from the bending stiffness *κ_b_* at a branch point between two fibers, we assume them to be equal *κ_b_* = *κ_f_* = *κ* because they both have contributions from the same underlying molecular crosslinking. This “equal constant” approximation reduces the number of free parameters in the model. In **Fig. S4F**, we consider the effects of having different values of buckling and bending stiffness.

By choosing the length scale as *l*_0_, force scale as *μ*, and time scale as *γl*_0_/*μ*, the dynamical equation (obtained by combining equation 1 with equations 2a–2c) can be rewritten in nondimensional form as

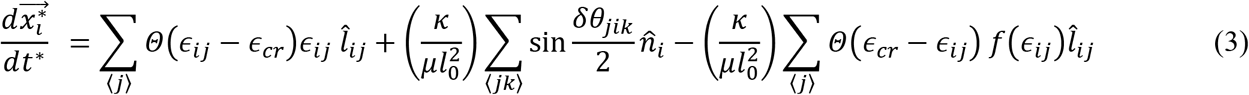

where starred quantities indicate dimensionless variables. The important nondimensional parameter that emerges is the ratio of bending to stretching stiffness, *κ*\* = *κ*/*μl*_0_^2^.

For Euler-Bernoulli beams, the stretching and bending moduli are given by *μ* = *EA* and *κ* = *EI*, where *E* is Young’s modulus and *A* and *I* are the cross-sectional area and the second moment of inertia, respectively (*35*). For a beam of circular cross-section, *A* = *πr*^2^, and *I* = *πr*^4^/4, and the expected bending to stretching ratio is then *κ*/(*μl*_0_^2^) = *r*^2^/(2*l*_0_^2^). A fibrin fiber is estimated to be *l*_0_ ≅ 10 μm long and 2*r* ≅ 280 nm thick (*22*),(*66*). Since fibrin is a bundle of protofibrils with bending modulus expected to be smaller than that of a uniform cylinder, we set the nondimensionalized ratio of bending to stretching stiffness parameter in the simulations to be *κ*\* = *k*·[*r*^2^/(2*l*_0_^2^)], where *r*^2^/(2*l*_0_^2^) is on the order of 10^-6^. Here, *k* is a factor less than unity, which accounts for the difference of fiber bending stiffness from the ideal limit of a uniform cylinder. The critical buckling strain is set to that of a uniform cylinder, *ϵ*_cr_ = −*π*^2^*r*^2^/*l*_0_^2^. The Young’s modulus of individual fibers was chosen to *E* = 1.4 MPa to match experimental shear modulus measurements. This comparatively low modulus is consistent with and lies within the wide range of measured stiffness of single fibrin fibers (*22*),(*23*),(*81*),(*82*). Using *E* = 1.4 MPa, we estimate the critical buckling load, *P_cr_* = *Eπ*^3^*r*^4^/*l*_0_^2^ ≅ 0.17 nN, which is about two orders of magnitude smaller than the contractile force of an activated platelet (~30 nN) (*51*). This suggests there is considerable platelet-induced fiber buckling in a typical PRP.

The network contraction due to platelets is modeled by prescribing shorter reference length for all bonds attached to a platelet. Assuming that an activated platelet can generate maximum contraction force *F_p_^max^*, which corresponds to maximum generated strain *ϵ_p_^max^* = *F_p_^max^/μ* the corresponding reference bond length is given by *<u>l_ij_</u>* / (1+*ϵ_p_^max^*). Thus, given values of platelet force correspond to the initial situation without any network deformation and externally applied shear strain. The forces in bonds attached to platelets are reduced via the initial relaxation at zero applied strain, and increase when shear strain is applied.

We perform the relaxation dynamics, and integrate equation (3) in time, until a mechanical equilibrium state is reached, implying that the change in net force becomes negligibly small (typically less than 10^-6^ %). Shear is applied on the network by displacing the upper and bottom boundary nodes by a fixed amount, while letting the other nodes move freely to balance forces in response to the applied strain. When the network reaches mechanical equilibrium, the applied strain is increased and the network relaxes to a new mechanical equilibrium. The increment of applied strain is chosen to be small (0.0005) to avoid large deformations near the boundary. Similar to the experiment, the upper and bottom boundaries were always clamped.

## Author contributions (alphabetically)

Conceptualization: KD, AG, SJL, AKR, AZ

Formal Analysis: KD, SJL, AKR, AZ

Methodology: KD, SJL, AKR, AZ

Investigation: MA, TC, KD, SJL, AKR, AZ

Visualization: KD, SJL, AKR, AZ

Supervision: KD, AKR

Writing—original draft: KD, AKR, AZ

Writing—review & editing: KD, AG, SJL, AKR, AZ

## Data and materials availability

All data needed to evaluate the conclusions in the paper are present in the paper and/or the Supplementary Materials.

## REFERENCES

1. J. W. Weisel, R. I. Litvinov, “Fibrin formation, structure and properties” in Fibrous Proteins: Structures and Mechanisms (Springer, Cham, 2017), Subcellular Biochemistry, pp. 405–456.

2. F. Burla, Y. Mulla, B. E. Vos, A. Aufderhorst-Roberts, G. H. Koenderink, From mechanical resilience to active material properties in biopolymer networks. Nature Reviews Physics. 1, 249–263 (2019).

3. T. A. E. Ahmed, E. V. Dare, M. Hincke, Fibrin: a versatile scaffold for tissue engineering applications. Tissue Eng. Part B Rev. 14, 199–215 (2008).

4. S. Kattula, J. R. Byrnes, A. S. Wolberg, Fibrinogen and Fibrin in Hemostasis and Thrombosis. Arterioscler. Thromb. Vasc. Biol. 37, e13–e21 (2017).

5. S. J. Pathare, W. Eng, S. J. J. Lee, A. K. Ramasubramanian, Fibrin prestress due to platelet aggregation and contraction increases clot stiffness. Biophysical Reports (2021) (available at https://www.sciencedirect.com/science/article/pii/S2667074721000227).

6. K. C. Gersh, C. Nagaswami, J. W. Weisel, Fibrin network structure and clot mechanical properties are altered by incorporation of erythrocytes. Thromb. Haemost. 102, 1169–1175 (2009).

7. T. Feller, S. D. A. Connell, R. A. S. Ariёns, Why Fibrin Biomechanical Properties Matter for Haemostasis and Thrombosis. J. Thromb. Haemost. (2021), doi:10.1111/jth.15531.

8. F. Pancaldi, O. V. Kim, J. W. Weisel, M. Alber, Z. Xu, Computational Biomechanical Modeling of Fibrin Networks and Platelet-Fiber Network Interactions. Current Opinion in Biomedical Engineering, 100369 (2022).

9. T. H. S. van Kempen, W. P. Donders, F. N. van de Vosse, G. W. M. Peters, A constitutive model for developing blood clots with various compositions and their nonlinear viscoelastic behavior. Biomech. Model. Mechanobiol. 15, 279–291 (2016).

10. A. S. G. van Oosten, X. Chen, L. Chin, K. Cruz, A. E. Patteson, K. Pogoda, V. B. Shenoy, P. A. Janmey, Emergence of tissue-like mechanics from fibrous networks confined by close-packed cells. Nature. 573, 96–101 (2019).

11. F. Ghezelbash, S. Liu, A. Shirazi-Adl, J. Li, Blood clot behaves as a poro-visco-elastic material. J. Mech. Behav. Biomed. Mater. 128, 105101 (2022).

12. B. Fereidoonnezhad, P. McGarry, A new constitutive model for permanent deformation of blood clots with application to simulation of aspiration thrombectomy. J. Biomech. 130, 110865 (2021).

13. C. Storm, J. J. Pastore, F. C. MacKintosh, T. C. Lubensky, P. A. Janmey, Nonlinear elasticity in biological gels. Nature. 435, 191–194 (2005).

14. N. Takeishi, T. Shigematsu, R. Enosaki, S. Ishida, S. Ii, S. Wada, Development of a mesoscopic framework spanning nanoscale protofibril dynamics to macro-scale fibrin clot formation. J. R. Soc. Interface. 18, 20210554 (2021).

15. S. Yesudasan, X. Wang, R. D. Averett, Fibrin polymerization simulation using a reactive dissipative particle dynamics method. Biomech. Model. Mechanobiol. 17, 1389–1403 (2018).

16. A. Zhmurov, O. Kononova, R. I. Litvinov, R. I. Dima, V. Barsegov, J. W. Weisel, Mechanical transition from α-helical coiled coils to β-sheets in fibrin(ogen). J. Am. Chem. Soc. 134, 20396–20402 (2012).

17. A. Mann, R. S. Sopher, S. Goren, O. Shelah, O. Tchaicheeyan, A. Lesman, Force chains in cell-cell mechanical communication. J. R. Soc. Interface. 16, 20190348 (2019).

18. M. Sarkar, J. Notbohm, Evolution of Force Chains Explains the Onset of Strain Stiffening in Fiber Networks. J. Appl. Mech., 1–19 (2022).

19. E. A. Ryan, L. F. Mockros, A. M. Stern, L. Lorand, Influence of a natural and a synthetic inhibitor of factor XIIIa on fibrin clot rheology. Biophys.J. 77, 2827–2836 (1999).

20. E. L. Hethershaw, A. L. Cilia La Corte, C. Duval, M. Ali, P. J. Grant, R. A. S. Ariëns, H. Philippou, The effect of blood coagulation factor XIII on fibrin clot structure and fibrinolysis. J. Thromb. Haemost. 12, 197–205 (2014).

21. D. C. Rijken, S. Abdul, J. J. M. C. Malfliet, F. W. G. Leebeek, S. Uitte de Willige, Compaction of fibrin clots reveals the antifibrinolytic effect of factor XIII. J. Thromb. Haemost. 14, 1453–1461 (2016).

22. J.-P. Collet, H. Shuman, R. E. Ledger, S. Lee, J. W. Weisel, The elasticity of an individual fibrin fiber in a clot. Proc. Natl. Acad. Sci. U. S. A. 102, 9133–9137 (2005).

23. W. Liu, C. R. Carlisle, E. A. Sparks, M. Guthold, The mechanical properties of single fibrin fibers. J. Thromb. Haemost. 8, 1030–1036 (2010).

24. J. Notbohm, A. Lesman, P. Rosakis, D. A. Tirrell, G. Ravichandran, Microbuckling of fibrin provides a mechanism for cell mechanosensing. J. R. Soc. Interface. 12, 20150320 (2015).

25. X. Xu, S. A. Safran, Nonlinearities of biopolymer gels increase the range of force transmission. Phys. Rev. E Stat. Nonlin. Soft Matter Phys. 92, 032728 (2015).

26. X. Ma, M. E. Schickel, M. D. Stevenson, A. L. Sarang-Sieminski, K. J. Gooch, S. N. Ghadiali, R. T. Hart, Fibers in the extracellular matrix enable long-range stress transmission between cells. Biophys. J. 104, 1410–1418 (2013).

27. S. Natan, Y. Koren, O. Shelah, S. Goren, A. Lesman, Long-range mechanical coupling of cells in 3D fibrin gels. Mol. Biol. Cell. 31, 1474–1485 (2020).

28. P. M. Nair, M. A. Meledeo, A. R. Wells, X. Wu, J. A. Bynum, K. P. Leung, B. Liu, A. Cheeniyil, A. K. Ramasubramanian, J. W. Weisel, A. P. Cap, Cold-stored platelets have better preserved contractile function in comparison with room temperature-stored platelets over 21 days. Transfusion. 61 Suppl 1, S68–S79 (2021).

29. Y. Sun, O. Oshinowo, D. R. Myers, W. A. Lam, A. Alexeev, Resolving the missing link between single platelet force and clot contractile force. iScience, 103690 (2021).

30. M. A. Panteleev, N. Korin, K. D. Reesink, D. L. Bark, J. M. E. M. Cosemans, E. E. Gardiner, P. H. Mangin, Wall shear rates in human and mouse arteries: Standardization of hemodynamics for in vitro blood flow assays: Communication from the ISTH SSC subcommittee on biorheology. J. Thromb. Haemost. 19, 588–595 (2021).

31. A. Sharma, A. J. Licup, K. A. Jansen, R. Rens, M. Sheinman, G. H. Koenderink, F. C. MacKintosh, Strain-controlled criticality governs the nonlinear mechanics of fibre networks. Nat. Phys. 12, 584–587 (2016).

32. J. C. Maxwell, L. On the calculation of the equilibrium and stiffness of frames. Lond. Edinb. Dublin Philos. Mag. J. Sci. 27, 294–299 (1864).

33. J. C. Maxwell, XLV. On reciprocal figures and diagrams of forces. The London, Edinburgh, and Dublin Philosophical Magazine and Journal of Science. 27, 250–261 (1864).

34. C. P. Broedersz, X. Mao, T. C. Lubensky, F. C. MacKintosh, Criticality and isostaticity in fibre networks. Nat. Phys. 7, 983–988 (2011).

35. L. D. Landau, E. M. Lifšic, E. M. Lifshitz, A. M. Kosevich, L. P. Pitaevskii, Theory of Elasticity: Volume 7 (Elsevier, 1986).

36. C. Heussinger, E. Frey, Floppy modes and nonaffine deformations in random fiber networks. Phys. Rev. Lett. 97, 105501 (2006).

37. P. R. Onck, T. Koeman, T. van Dillen, E. van der Giessen, Alternative explanation of stiffening in cross-linked semiflexible networks. Phys. Rev. Lett. 95, 178102 (2005).

38. J. L. Shivers, J. Feng, A. S. G. van Oosten, H. Levine, P. A. Janmey, F. C. MacKintosh, Compression stiffening of fibrous networks with stiff inclusions. Proc. Natl. Acad. Sci. U. S. A. 117, 21037–21044 (2020).

39. K. F. Freund, K. P. Doshi, S. L. Gaul, D. A. Claremon, D. C. Remy, J. J. Baldwin, S. M. Pitzenberger, A. M. Stern, Transglutaminase inhibition by 2-[(2-oxopropyl)thio]imidazolium derivatives: mechanism of factor XIIIa inactivation. Biochemistry. 33, 10109–10119 (1994).

40. M. Bathe, C. Heussinger, M. M. A. E. Claessens, A. R. Bausch, E. Frey, Cytoskeletal bundle mechanics. Biophys. J. 94, 2955–2964 (2008).

41. S. Britton, O. Kim, F. Pancaldi, Z. Xu, R. I. Litvinov, J. W. Weisel, M. Alber, Contribution of nascent cohesive fiber-fiber interactions to the non-linear elasticity of fibrin networks under tensile load. Acta Biomater. 94, 514–523 (2019).

42. J. Xia, L.-H. Cai, H. Wu, F. C. MacKintosh, D. A. Weitz, Anomalous mechanics of Zn2+-modified fibrin networks. Proc. Natl. Acad. Sci. U. S. A. 118 (2021), doi:10.1073/pnas.2020541118.

43. I. K. Piechocka, R. G. Bacabac, M. Potters, F. C. Mackintosh, G. H. Koenderink, Structural hierarchy governs fibrin gel mechanics. Biophys. J. 98, 2281–2289 (2010).

44. C. P. Broedersz, F. C. MacKintosh, Molecular motors stiffen non-affine semiflexible polymer networks. Soft Matter. 7, 3186–3191 (2011).

45. A. Sharma, A. J. Licup, R. Rens, M. Vahabi, K. A. Jansen, G. H. Koenderink, F. C. MacKintosh, Strain-driven criticality underlies nonlinear mechanics of fibrous networks. Physical review.E, Statistical, nonlinear, and soft matter physics. 94 (2016).

46. S. Münster, L. M. Jawerth, B. A. Leslie, J. I. Weitz, B. Fabry, D. A. Weitz, Strain history dependence of the nonlinear stress response of fibrin and collagen networks. Proc. Natl. Acad. Sci. U. S. A. 110, 12197–12202 (2013).

47. J. Feng, H. Levine, X. Mao, L. M. Sander, Nonlinear elasticity of disordered fiber networks. Soft Matter. 12, 1419–1424 (2016).

48. E. Conti, F. C. Mackintosh, Cross-linked networks of stiff filaments exhibit negative normal stress. Phys. Rev. Lett. 102, 088102 (2009).

49. M. Bouzid, E. Del Gado, Network Topology in Soft Gels: Hardening and Softening Materials. Langmuir. 34, 773–781 (2018).

50. J. Hanke, D. Probst, A. Zemel, U. S. Schwarz, S. Köster, Dynamics of force generation by spreading platelets. Soft Matter. 14, 6571–6581 (2018).

51. W. A. Lam, O. Chaudhuri, A. Crow, K. D. Webster, T.-D. Li, A. Kita, J. Huang, D. A. Fletcher, Mechanics and contraction dynamics of single platelets and implications for clot stiffening. Nat. Mater. 10, 61–66 (2011).

52. K. F. Standeven, A. M. Carter, P. J. Grant, J. W. Weisel, I. Chernysh, L. Masova, S. T. Lord, R. A. S. Ariëns, Functional analysis of fibrin {gamma}-chain cross-linking by activated factor XIII: determination of a cross-linking pattern that maximizes clot stiffness. Blood. 110, 902–907 (2007).

53. C. Duval, P. Allan, S. D. A. Connell, V. C. Ridger, H. Philippou, R. A. S. Ariëns, Roles of fibrin α-and γ-chain specific cross-linking by FXIIIa in fibrin structure and function. Thromb. Haemost. 111, 842–850 (2014).

54. Y. Qiu, A. C. Brown, D. R. Myers, Y. Sakurai, R. G. Mannino, R. Tran, B. Ahn, E. T. Hardy, M. F. Kee, S. Kumar, G. Bao, T. H. Barker, W. A. Lam, Platelet mechanosensing of substrate stiffness during clot formation mediates adhesion, spreading, and activation. Proc. Natl. Acad. Sci. U. S. A. 111, 14430–14435 (2014).

55. P. A. Janmey, J. P. Winer, J. W. Weisel, Fibrin gels and their clinical and bioengineering applications. J. R. Soc. Interface. 6, 1–10 (2009).

56. J. V. Shah, P. A. Janmey, Strain hardening of fibrin gels and plasma clots. Rheola Acta. 36, 262–268 (1997).

57. N. A. Kurniawan, J. Grimbergen, J. Koopman, G. H. Koenderink, Factor XIII stiffens fibrin clots by causing fiber compaction. J. Thromb. Haemost. 12, 1687–1696 (2014).

58. A. Kabla, L. Mahadevan, Nonlinear mechanics of soft fibrous networks. J. R. Soc. Interface. 4, 99–106 (2007).

59. F. Meng, E. M. Terentjev, Theory of semiflexible filaments and networks. Polymers (Basel). 9, 52 (2017).

60. M. L. Gardel, J. H. Shin, F. C. MacKintosh, L. Mahadevan, P. Matsudaira, D. A. Weitz, Elastic behavior of cross-linked and bundled actin networks. Science. 304, 1301–1305 (2004).

61. R. C. Picu, Mechanics of random fiber networks—a review. Soft Matter. 7, 6768–6785 (2011).

62. F. Maksudov, A. Daraei, A. Sesha, K. A. Marx, M. Guthold, V. Barsegov, Strength, Deformability and Toughness of Uncrosslinked Fibrin Fibers from Theoretical Reconstruction of Stress-Strain Curves. Acta Biomater. (2021), doi:10.1016/j.actbio.2021.09.050.

63. O. V. Kim, R. I. Litvinov, J. W. Weisel, M. S. Alber, Structural basis for the nonlinear mechanics of fibrin networks under compression. Biomaterials. 35, 6739–6749 (2014).

64. X. Liang, I. Chernysh, P. K. Purohit, J. W. Weisel, Phase transitions during compression and decompression of clots from platelet-poor plasma, platelet-rich plasma and whole blood. Acta Biomater. 60, 275–290 (2017).

65. U. Windberger, V. Glanz, L. Ploszczanski, Laboratory Rat Thrombi Lose One-Third of Their Stiffness When Exposed to Large Oscillating Shear Stress Amplitudes: Contrasting Behavior to Human Clots. International Journal of Translational Medicine. 2, 332–344 (2022).

66. S.-J. J. Lee, D. M. Nguyen, H. S. Grewal, C. Puligundla, A. K. Saha, P. M. Nair, A. P. Cap, A. K. Ramasubramanian, Image-based analysis and simulation of the effect of platelet storage temperature on clot mechanics under uniaxial strain. Biomech. Model. Mechanobiol. 19, 173–187 (2019).

67. M. Alzweighi, R. Mansour, J. Lahti, U. Hirn, A. Kulachenko, The influence of structural variations on the constitutive response and strain variations in thin fibrous materials. Acta Mater. 203, 116460 (2021).

68. C. Heussinger, E. Frey, Force distributions and force chains in random stiff fiber networks. Eur. Phys. J. E Soft Matter. 24, 47–53 (2007).

69. M. S. Rudnicki, H. A. Cirka, M. Aghvami, E. A. Sander, Q. Wen, K. L. Billiar, Nonlinear strain stiffening is not sufficient to explain how far cells can feel on fibrous protein gels. Biophys. J. 105, 11–20 (2013).

70. A. S. Abhilash, B. M. Baker, B. Trappmann, C. S. Chen, V. B. Shenoy, Remodeling of fibrous extracellular matrices by contractile cells: predictions from discrete fiber network simulations. Biophys. J. 107, 1829–1840 (2014).

71. O. V. Kim, Z. Xu, E. D. Rosen, M. S. Alber, Fibrin networks regulate protein transport during thrombus development. PLoS Comput. Biol. 9, e1003095 (2013).

72. R. S. Voronov, T. J. Stalker, L. F. Brass, S. L. Diamond, Simulation of intrathrombus fluid and solute transport using in vivo clot structures with single platelet resolution. Ann. Biomed. Eng. 41, 1297–1307 (2013).

73. A. V. Belyaev, M. A. Panteleev, F. I. Ataullakhanov, Threshold of microvascular occlusion: injury size defines the thrombosis scenario. Biophys. J. 109, 450–456 (2015).

74. V. Govindarajan, S. Zhu, R. Li, Y. Lu, S. L. Diamond, J. Reifman, A. Y. Mitrophanov, Impact of tissue factor localization on blood clot structure and resistance under venous shear. Biophys. J. 114, 978–991 (2018).

75. Y. L. Han, P. Ronceray, G. Xu, A. Malandrino, R. D. Kamm, M. Lenz, C. P. Broedersz, M. Guo, Cell contraction induces long-ranged stress stiffening in the extracellular matrix. Proc. Natl. Acad. Sci. U. S. A. 115, 4075–4080 (2018).

76. N. A. Kurniawan, T. H. S. van Kempen, S. Sonneveld, T. T. Rosalina, B. E. Vos, K. A. Jansen, G. W. M. Peters, F. N. van de Vosse, G. H. Koenderink, Buffers strongly modulate fibrin self-assembly into fibrous networks. Langmuir. 33, 6342–6352 (2017).

77. C. J. Jen, L. V. McIntire, The structural properties and contractile force of a clot. Cell Motil. 2, 445–455 (1982).

78. A. S. Caroline, S. R. Wayne, W. E. Kevin, NIH Image to ImageJ: 25 years of image analysis. Nat. Methods. 9, 671 (2012).

79. C. Geuzaine, J.-F. Remacle, Gmsh: A 3-D finite element mesh generator with built-in pre-and post-processing facilities. Int. J. Numer. Methods Eng. 79, 1309–1331 (2009).

80. S. Timoshenko, Theory of elastic stability (Tata McGraw-Hill Education, ed. 2, 1963).

81. J. R. Houser, N. E. Hudson, L. Ping, E. T. O’Brien 3rd, R. Superfine, S. T. Lord, M. R. Falvo, Evidence that αC region is origin of low modulus, high extensibility, and strain stiffening in fibrin fibers. Biophys. J. 99, 3038–3047 (2010).

82. M. Guthold, W. Liu, E. A. Sparks, L. M. Jawerth, L. Peng, M. Falvo, R. Superfine, R. R. Hantgan, S. T. Lord, A comparison of the mechanical and structural properties of fibrin fibers with other protein fibers. Cell Biochem. Biophys. 49, 165–181 (2007).

