## Supplementary Information for "Clots reveal anomalous elastic behavior of fiber networks"

Zakharov *et al.*

**This PDF file includes:**

Figs. S1 to S5

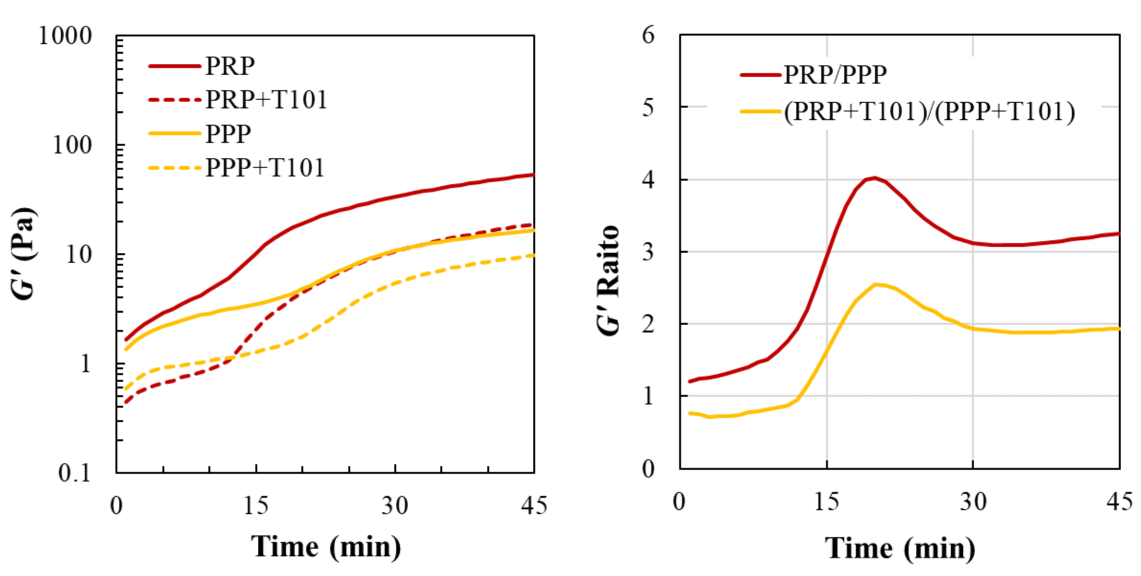


**Fig. S1. Gelation of PPP and PRP clots**

**(A)** Representative gelation profiles for platelet poor plasma (PPP) and platelet rich plasma (PRP) clots, showing elastic storage moduli (*G′*) vs. time for clots prepared with and without T101 as a crosslinking inhibitor. **(B)** After ~30 min, the average relative increase in *G′* associated with the presence of platelets (i.e., ratio of PRP/PPP) was approximately threefold for crosslinked fibrin clots and approximately twofold for the uncrosslinked (T101-treated) case.

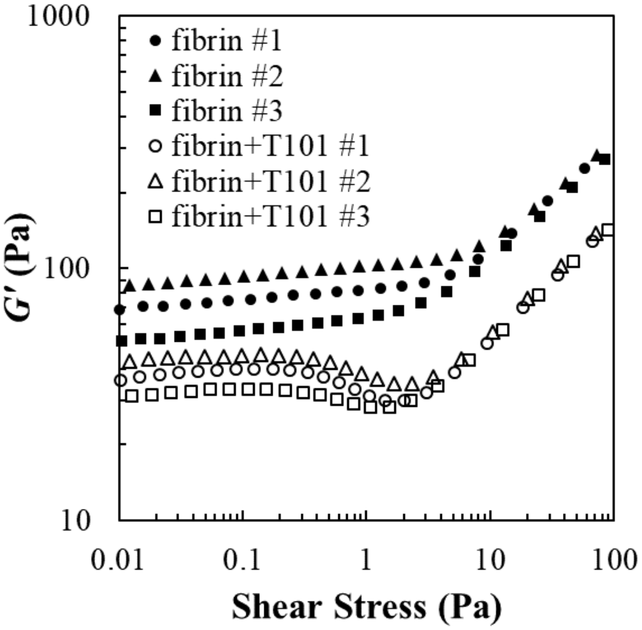


**Fig. S2. Shear rheology of pure fibrin networks**

Elastic storage modulus *G′* vs. shear stress for fibrin gels, comparing crosslinked and uncrosslinked fibrin, showing $n = 3$ identically prepared replicates for each case. Fibrin gels were formed using 3 mg/mL fibrinogen with 1 U/mL thrombin and 20 mM $CaCl_2$. For the uncrosslinked gels, 100 μM T101 was added as an inhibitor. Both types of fibrin gels are stress-stiffening. However the uncrosslinked case exhibits a dip near 1 Pa, which is similar to the model-predicted behavior and T101-treated PPP data in the main text (**Fig. 2**).

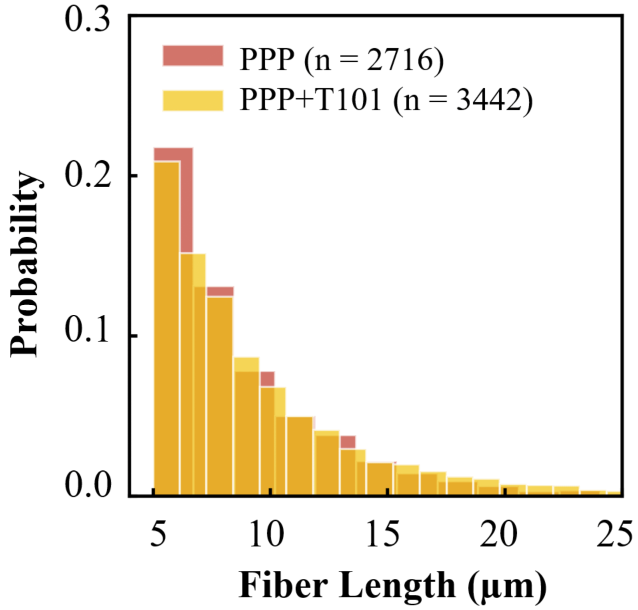


**Fig. S3. Distribution of fiber lengths in crosslinked and uncrosslinked PPP clots.**
Fiber lengths for PPP clots with and without T101 were determined from a set of fluorescence microscopy images at equal magnification (3 images for each case). A threshold of 5 µm was used to filter artifactually short segments and image fragments. The histograms reveal that there is no substantial difference in the distribution of fiber lengths between crosslinked and uncrosslinked fibrin.

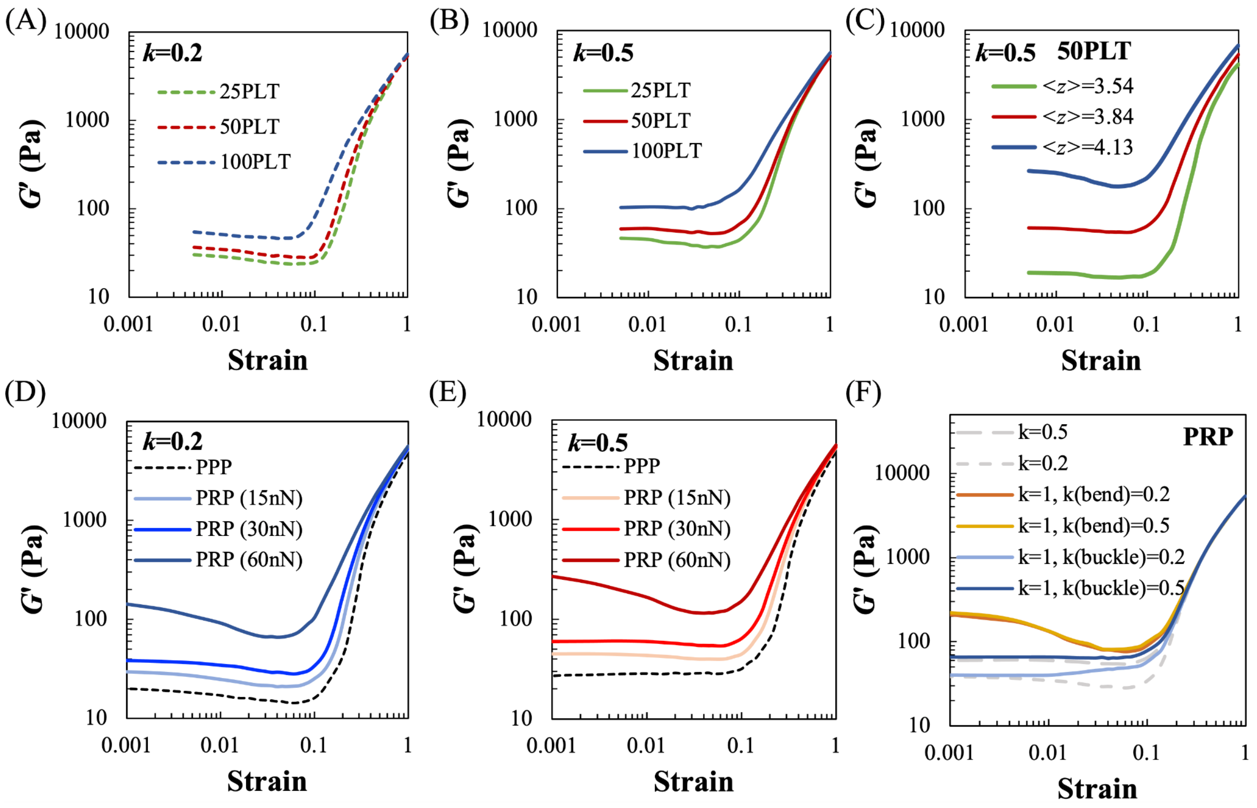


**Fig. S4. Effect of number of platelet aggregates, connectivity, platelet force and fiber stiffness on the response of active networks.**

The critical strain at which stiffening transition takes place strongly depends on the number of platelets. Simulations reveal that increasing platelet density leads to lowering critical strain in both uncrosslinked **(A)** and crosslinked **(B)** clots. Contracting platelet aggregates pull out available soft bending and buckling modes, and thus the transition to the stiff stretching dominated mode occurs at smaller strains. Platelet aggregates prestress the network and create additional buckling, which results in softening of the network, and this effect is more pronounced in uncrosslinked clots at lower $k$, or in over-coordinated networks at larger $z$ **(C)**, because bending is limited in networks at larger $z$ and platelet aggregates cause more fibers to buckle. Increasing platelet contractile force stiffen the network in a non-linear manner **(D, E)**. Reducing exclusively bending or buckling stiffness **(F)** shows that platelets significantly stiffen the networks with smaller bending resistance at small strains (orange and red lines in **(F)**). Since bending is energetically cheaper (easy to change the angle between fibers at branch points), platelets can efficiently form force chains without fiber buckling, along which the network is reinforced. With strain, buckling becomes unavoidable, this results in network softening until transition to stiff stretching mode. Platelets in networks with rigid branch points (blue and light blue lines in **(F)**), conversely, pull out available energetically cheap buckling modes and demonstrate only bending dominated regime (the plateau in *G*'). Since bending is energetically unfavorable, platelets deform the network more uniformly and at small ranges, and thus the network is weakly stiffened by platelets.

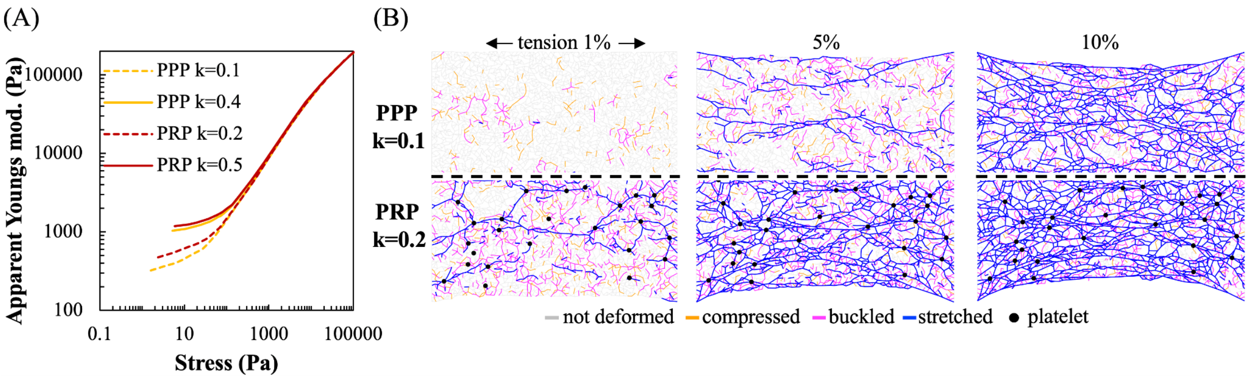


**Fig. S5. Behavior of active and passive networks under uniaxial tension.**

Simulated clots under uniaxial tension also demonstrate stiffening transitions associated with an increasing number of stretched fibers in the network. Similar to shear simulation results, the transition is predicted to occur at lower stresses in uncrosslinked clots, whereas platelet aggregates are expected to increase the critical stress **(A)**. Platelet aggregates actively contract the network in PRP clots and form force chains of stretched fibers even at low strains (**B**, PRP 1% strain), which increase in number at larger applied loading. Even though the applied stress was uniform at the left and right edges, the stretching is localized along a few fibers in under-coordinated networks (**B**, PPP 5% strain). The effect of platelets eliminates at larger strains when applied force significantly exceeds the contractile force by platelets. In this regime, both PPP and PRP clots have a similar number of stretched fibers (B, 10% strain) and the same stiffness **(A)**.
